# Adapterama III: Quadruple-indexed, double/triple-enzyme RADseq libraries (2RAD/3RAD)

**DOI:** 10.1101/205799

**Authors:** Natalia J Bayona-Vásquez, Travis C Glenn, Troy J Kieran, Todd W Pierson, Sandra L Hoffberg, Peter A Scott, Kerin E Bentley, John W Finger, Swarnali Louha, Nicholas Troendle, Píndaro Díaz-Jaimes, Rodney Mauricio, Brant C Faircloth

**Author notes:** Department of Ecology and Evolutionary Biology, University of Tennessee, Knoxville, TN 37996, USA. Department of Ecology, Evolution and Environmental Biology, Columbia University, New York, NY 10027, USA. Department of Ecology and Evolutionary Biology, University of California, Los Angeles, CA 90095, USA. LeafWorks Inc., 125 South Main Street #150, Sebastopol, CA 95472, USA. Department of Biological Sciences, Auburn University, Auburn, AL 36849, USA. Department of Natural, Health, and Mathematical Sciences, MidAmerica Nazarene University, 2030 E. College Way, Olathe, KS 66062, USA. equal contributions as first authors (Natalia J Bayona-Vásquez, Travis C Glenn). Corresponding Authors: Natalia Bayona-Vásquez Dept. of EHS, Environmental Health Science Bldg., University of Georgia, Athens, GA 30602, USA Email address, Travis Glenn Dept. of EHS, Environmental Health Science Bldg., University of Georgia, Athens, GA 30602, USA.

## Abstract

Molecular ecologists frequently use genome reduction strategies that rely upon restriction enzyme digestion of genomic DNA to sample consistent portions of the genome from many individuals (e.g., RADseq, GBS). However, researchers often find the existing methods expensive to initiate and/or difficult to implement consistently, especially due to the inability to highly-multiplex samples to fill entire sequencing lanes. Here, we introduce a low-cost and highly robust approach for the construction of dual-digest RADseq libraries that relies on adapters and primers designed in *Adapterama I*. Major features of our method include: 1) minimizing the number of processing steps; 2) focusing on a single strand of sample DNA for library construction, allowing the use of a non-phosphorylated adapter on one end; 3) ligating adapters in the presence of active restriction enzymes, thereby reducing chimeras; 4) including an optional third restriction enzyme to cut apart adapter-dimers formed by the phosphorylated adapter, thus increasing the efficiency of adapter ligation to sample DNA, which is particularly effective when only low quantity/quality DNA samples are available; 5) interchangeable adapter designs; 6) incorporating variable-length internal indexes within the adapters to increase the scope of sample indexing, facilitate pooling, and increase sequence diversity; 7) maintaining compatibility with universal dual-indexed primers and thus, Illumina sequencing reagents and libraries; and, 8) easy modification for the identification of PCR duplicates. We present eight adapter designs that work with 72 restriction enzyme combinations. We demonstrate the efficiency of our approach by comparing it with existing methods, and we validate its utility through the discovery of many variable loci in a variety of non-model organisms. Our 2RAD/3RAD method is easy to perform, has low startup costs, has increased utility with low-concentration input DNA, and produces libraries that can be highly-multiplexed and pooled with other Illumina libraries.

## Introduction

Although next-generation DNA sequencing (NGS) facilitates data collection at low cost, it is not yet economically or computationally feasible for most ecological projects to sequence whole genomes from many individuals or from organisms with large genomes. However, many questions can be addressed with a small fraction of the genome (DeWoody & DeWoody, 2005; Cariou, Duret, & Charlat, 2013; Pante et al., 2015; Andrews et al., 2016). Thus, researchers have created a variety of strategies to sample a consistent portion of the genome from large numbers of individuals at low cost (Harvey et al., 2016; Heyduk et al., 2016; Glenn & Faircloth, 2016). One of the most popular genome sampling strategies uses restriction enzymes (REs) to reduce genome complexity and sequence a set of orthologous loci across individuals (Restriction site Associated DNA sequencing, RADseq; Miller et al., 2007; Baird et al., 2008). Key advantages of the RADseq approach include: 1) the lack of a need for a reference genome (although it is better to have one [e.g., Shafer et al., 2016]); 2) the relatively low cost of library preparations; 3) the applicability of techniques with minimal modification across a broad spectrum of organisms; and 4) the availability of suitable software for data analyses (e.g., Stacks; Catchen et al., 2011; Catchen et al., 2013; pyRAD; Eaton, 2014).

Many variants of the general RADseq approach have been developed (Andrews et al. 2016; Salas-Linaza & Oono, 2018), including but not limited to: the original method (RAD; Miller et al., 2007; Baird et al., 2008), genotype-by-sequencing (GBS; Elshire et al., 2011), 2-enzyme GBS (Poland & Rife, 2012), dual-digest RADseq (ddRAD; Peterson et al., 2012), 2bRAD (Wang et al., 2012), ezRAD (Toonen et al., 2013), and PCR duplicate removal (quaddRAD; Franchini et al., 2017). Although some of these RADseq approaches are in widespread use, they also have well-documented limitations (Davey et al., 2011; Andrews et al., 2014) which have been thoroughly reviewed in other publications (Andrews et al., 2016; Harvey et al., 2016; Heyduk et al., 2016).

Limitations of methods that pool DNA samples from multiple individuals within putative populations are particularly acute (Andrews et al., 2016); thus, we focus on methods where individual samples are indexed such that individual organisms can be genotyped. Limitations of current individual-based RADseq methods include: i) high up-front costs for adapters (∼$4,550 USD, for 12 pools of 48 samples; Peterson et al., 2012; Salas-Linaza & Oono, 2018), which limits experimental flexibility; ii) that adapters are phosphorylated in both ends, and thus, can form adapter dimers; iii) the use of an optional step to incorporate a biotin-containing primer and streptavidin beads to separate correct constructs; iv) the inability to reduce chimera formation; v) the fact that no sequence diversity is present across the restriction recognition sequence of the resulting libraries, which constrains sequencing options; vi) the need for moderate to high amounts of high-molecular-weight DNA; vii) limited ability to multiplex high numbers of libraries, and thus, high sequencing costs; and viii) workflows of varying complexity. Thus, developing RADseq methods that reduce the cost of adapters and primers, vastly improve sample multiplexing, reduce the amount of input DNA, improve flexibility, increase consistency, and simplify the workflow would be helpful to many researchers.

Here, we present an alternative approach (2RAD/3RAD) for preparing RADseq libraries, which is similar in spirit to and builds upon the strengths of ddRAD, while also addressing each of the limitations summarized above and described below in our Methodological Objectives. In brief, 2RAD/3RAD uses DNA that is digested with multiple REs and ligates adapters in the presence of functional REs, followed by PCR with primers developed in *Adapterama I* (Glenn et al., 2016) to make fully active quadruple-indexed Illumina libraries that can be highly-multiplexed (Figs. 1 and S1). Although some of our adapter designs and working procedures have been implemented and published during the development of our method (e.g., Graham et al., 2015; Hoffberg et al., 2016; Scott, Glenn, & Rissler, 2017), additional designs, design details, flexibility, advantages and disadvantages of the system have yet to be described. Below, we explain the design goals and rationale for our approach, detail how we have implemented the method, demonstrate that large numbers of polymorphic loci can be discovered from a broad array of organisms using just one of the possible variations of our method, and discuss these results and additional work to extend this approach.

**Figure 1.**
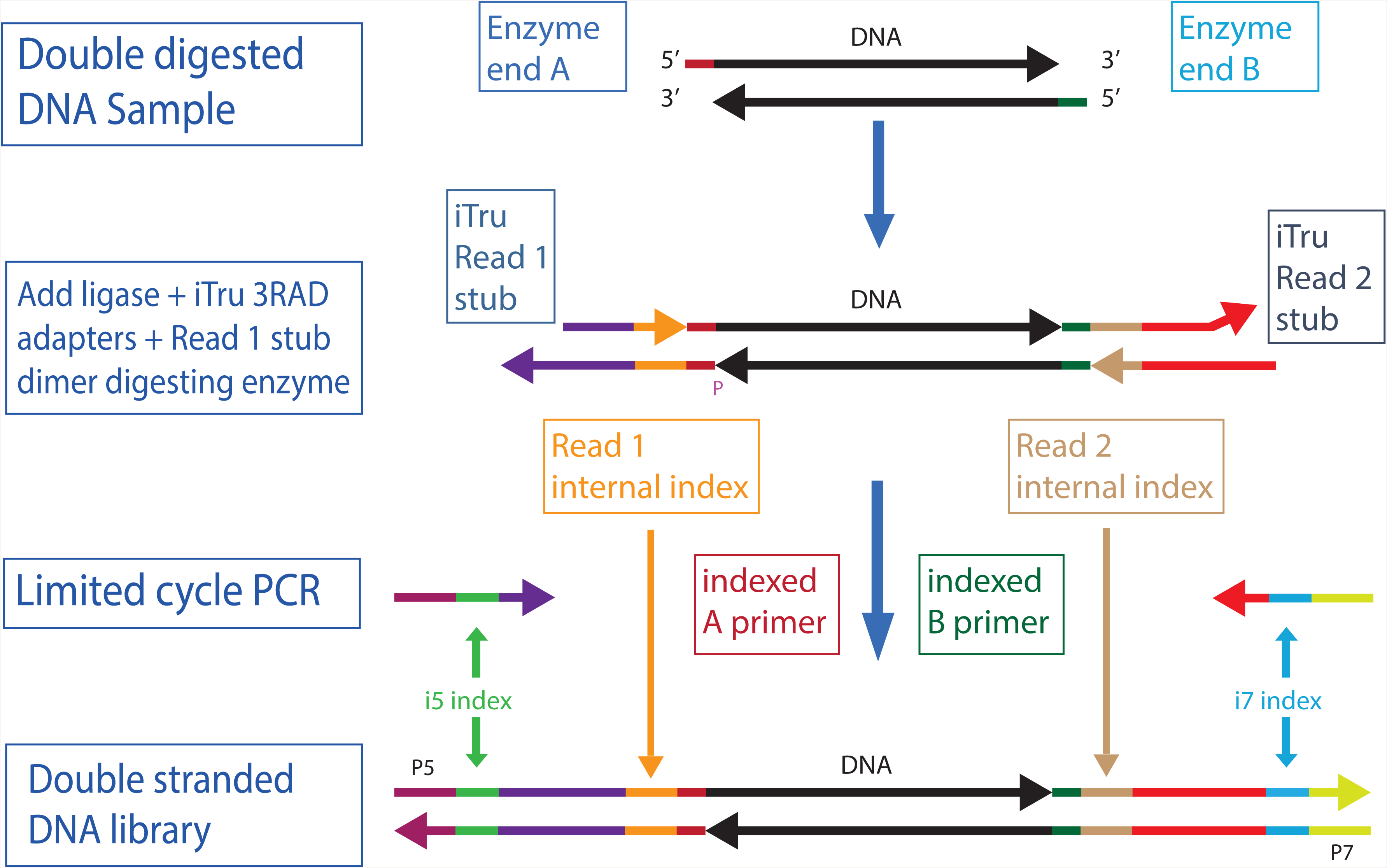
Overview 3RAD library construction. Genomic DNA is digested with two REs (A and B). Adapters are ligated to the digested DNA, but only the bottom strand has functional adapters. The top strand has shorter, non-functional versions of the adapters. The ligation products are then used in a limited cycle PCR with iTru5 and iTru7 primers to form fully active double-stranded DNA molecules. The color-scheme follows those of Glenn *et al.* (2016) and Hoffberg *et al.* (2016). **Note: These will be added individually to PeerJ with each file upload. *Don’t* include ‘Figure 1‘; just add the title and description separately. Titles are in bold and descriptions are in plain font.**

## Materials & Methods

### Methodological objectives

Our overall goal was to develop a ddRAD-style method with the following characteristics: 1) a simplified workflow with few consecutive buffer exchanges; 2) sequential or simultaneous digestion of DNA and ligation of adapters; 3) reduced chimera formation; 4) increased library efficiency through suppressed adapter dimer formation; 5) hierarchical combinatorial indexing to facilitate efficient multiplexing of many samples; 6) reduced costs, both for initial buy-in (i.e., cost of all reagents to start using the method) and per sample prepared; and 7) facilitation of pooling with any other Illumina library type. We built upon the adapter design and methods of Glenn & Schable (2005) to achieve the first four goals, whereas we extended the work of Faircloth & Glenn (2012) and Glenn et al. (2016) to achieve the last three goals (Files S1-S2).

### Methodological approach

We achieved our first three design goals by using reagents that allow simultaneous digestion of the sample DNA and ligation of the adapters onto the sample DNA (Glenn & Schable, 2005). Simultaneous digestion of DNA with multiple REs requires that the enzymes are active in the same buffer and at the same temperature. New England Biolabs (NEB; Ipswich, MA, USA) has developed many enzymes that retain high activity in a single buffer (CutSmart) and describes the activity of their enzymes in their other standard buffer formulations (Table 1). Because T4 DNA ligase can be used in the same buffers as most REs, if the buffers are supplemented with ATP, researchers can start by digesting DNA, then add ligase and ATP to the digestion reaction and change the temperature to promote ligation. By cycling between temperatures that promote ligation, then digestion, multiple times, reactions can be driven to highly efficient outcomes (i.e., high proportions of the input DNA will be cut and will have adapters ligated onto the ends; Figs. S1–S2). A major distinction between the methods of Glenn and Schable (2005) and those needed here is that a single blunt-ended 5’ phosphorylated adapter (Super SNX) was used in the prior work, resulting in identical adapters on each end of the resulting libraries, whereas Illumina libraries require unique adapter sequences on each end of the library molecules (Fig. 1). By focusing on a single strand of the template, rather than both strands, it is possible to use only one adapter that is phosphorylated (i.e., Read 1 adapter, bottom strand; Figs. 1, S1-S2) in 2RAD/3RAD, whereas the other adapter can use plain oligonucleotides for both strands (File S3), forming fewer adapter chimeras.

**Table 1.** Enzyme combinations and characteristics. Four design sets each for Read1 (R1) and Read2 (R2) are given. For 2RAD, any two enzymes combination of Read 1 and Read 2 in black can be used. For 3RAD, the third enzyme (in blue) blocks adapter-dimer formation of the Read 1 adapter (Supplemental File S3). Digestion efficiency is given for three NEB buffers (2.1, 3.1, and CutSmart^®^), with the best conditions highlighted in green, and poor or important non-standard conditions in red. Sensitivity to methylation in the template sequence is given, as is the optimal temperature for digestion and the number of bases in the recognition sequence. Note: Some REs are available as high-fidelity (HF, i.e.: NheI-HF, SpeI-HF, and NsiI-HF), all these have 100% efficiency on CutSmart^®^ Buffer.

| Read 1 Adapter Sets |  |  |  |  |  |  |  | Read 2 Adapter Sets |  |  |  |  |  |  |  |
| --- | --- | --- | --- | --- | --- | --- | --- | --- | --- | --- | --- | --- | --- | --- | --- |
| Set | Enzyme | NEB Buffer |  |  | CpG meth | Cut Temp | Base Cutter | Set | Enzyme | NEB Buffer |  |  | CpG meth | Cut Temp | Base Cutter |
|  |  | 2.1 | 3.1 | CutSmart® |  |  |  |  |  | 2.1 | 3.1 | CutSmart® |  |  |  |
| R1.A | NheI | 100 | 10 | 100 | +/- | 37 | 6 | R2.1 | EcoRI-HF | 100 | 10 | 100 | +/- | 37 | 6 |
|  | XbaI | 100 | 75 | 100 | - | 37 | 6 |  | MfeI-HF | 25 | 10 | 100 | - | 37 | 6 |
|  | SpeI | 100 | 25 | 100 | - | 37 | 6 |  | ApoI | 75 | 100 | 75 | - | 50* | 6 |
| R1.B | ClaI | 50 | 50 | 100 | + | 37 | 6 | R2.2 | BamHI-HF | 50 | 10 | 100 | - | 37 | 6 |
|  | MspI | 100 | 50 | 100 | - | 37 | 4 |  | BclI | 100 | 100 | 75 | - | 50* | 6 |
|  | TaqαI | 75 | 100 | 100 | - | 65 | 4 |  | BstYI | 100 | 75 | 100 | - | 60** | 6 |
| R1.C | PstI-HF | 75 | 50 | 100 | - | 37 | 6 | R2.3 | DdeI | 100 | 100 | 100 | - | 37 | 4 |
|  | PstI | 75 | 100 | 50 | - | 37 | 6 |  |  |  |  |  |  |  |  |
|  | NsiI | 75 | 100 | 25 | - | 37 | 6 |  |  |  |  |  |  |  |  |
| R1.D | CviQI | 100 | 100 | 75 | - | 37 | 6 | R2.4 | HindII-HF | 100 | 10 | 100 | - | 37 | 6 |
|  | NdeI | 100 | 100 | 100 | - | 37 | 6 |  | HindIII | 100 | 50 | 50 | - | 37 | 6 |
|  | MseI | 100 | 75 | 100 | - | 37 | 4 |  |  |  |  |  |  |  |  |
|  | AseI | 50 | 100 | 10 | - | 37 | 6 |  |  |  |  |  |  |  |  |
|  | BfaI | 10 | 10 | 100 | - | 37 | 4 |  |  |  |  |  |  |  |  |

We achieved our fourth design goal through the design of our adapters. During ligation, double-stranded adapters that are modified versions of the TruSeq Read 1 and Read 2 sequences (Table 2) are ligated onto each fragment of DNA. Oligonucleotide sequences for all versions of all adapter designs are given in File S3. We ordered these in plates from Integrated DNA Technologies (IDT; Coralville, IA, USA) and prepared them following directions in File S4. The iTru Read 2 adapters are unphosphorylated on the 5’ end and will not self-ligate to form dimers. The iTru Read 1 adapter is a perfect match to the sticky end of the insert DNA, but the adapter does not have the correct bases to recreate the restriction site used to cut the sample DNA. Because the iTru Read 1 adapter is phosphorylated on the 5’ end, it can and will form dimers.

**Table 2.**
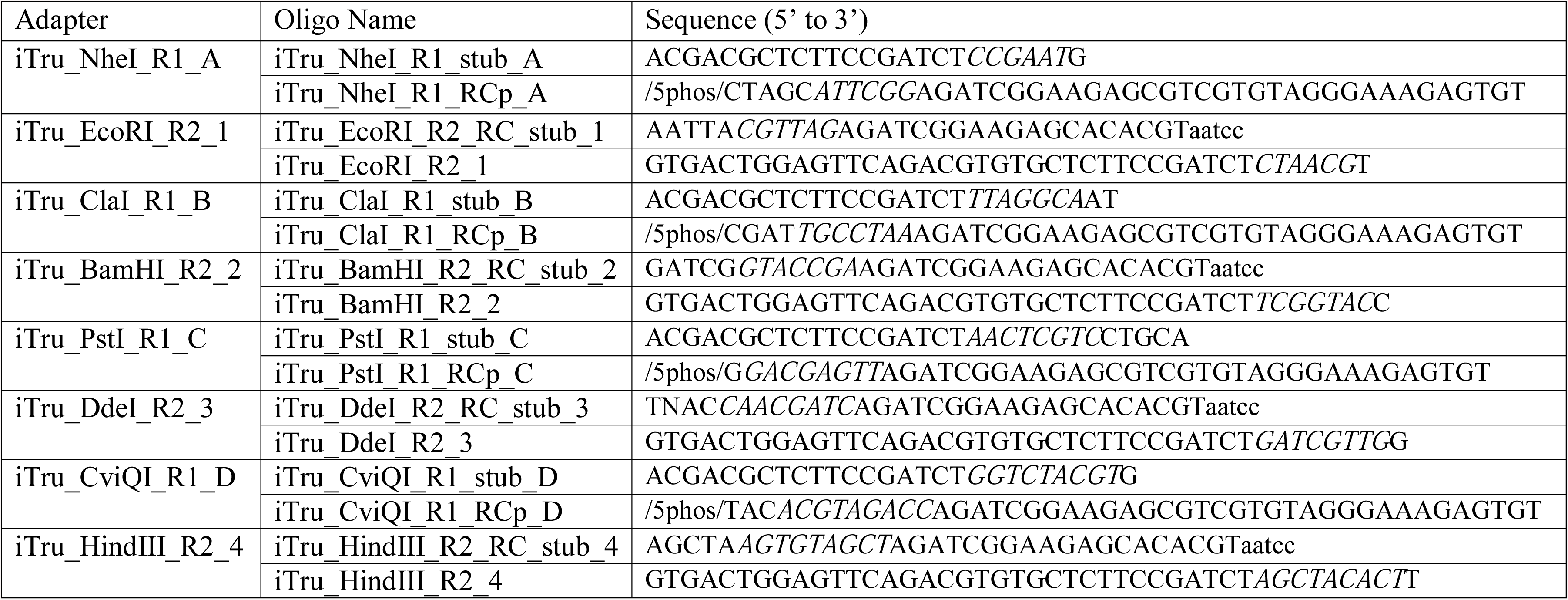
Example 2RAD/3RAD adapter stub sequences. Groups of 4 adapters form a balanced set, all eight complete sets are available in Appendix S3. Non-complementary sequences are given in lower case. Tag sequences are in italics. Adapters must be hydrated and annealed prior to use (Appendix S4).

For 2RAD, we selected sets of low-cost, type II REs that form unique cohesive-ends (i.e., incompatible sticky-ends). For 3RAD, we further achieved our fourth design goal by using these two REs (e.g., XbaI and EcoRI; Table 1) with a third RE (e.g., NheI) that produces a cohesive-end compatible with one of the other REs (e.g., XbaI; Fig. 2). We then assigned the two REs with compatible cohesive-ends (e.g., XbaI, and NheI) to Illumina Read 1 adapter stub sequences (Glenn et al., 2016) and assigned the incompatible REs (e.g., EcoRI) to Read 2 adapter stubs. Next, we designed the Read 1 stubs such that if they self-ligated to form Read 1 adapter-dimers, they create the recognition sequence for the third RE (e.g., NheI; File S3; Glenn & Schable, 2005). Similarly, Read 1 adapters ligated to genomic DNA with third RE cut-sites will recreate the recognition sequence for the third RE. See Fig. 2 for a graphical representation of one example of this design using the REs XbaI, EcoRI, and NheI.

**Figure 2.**
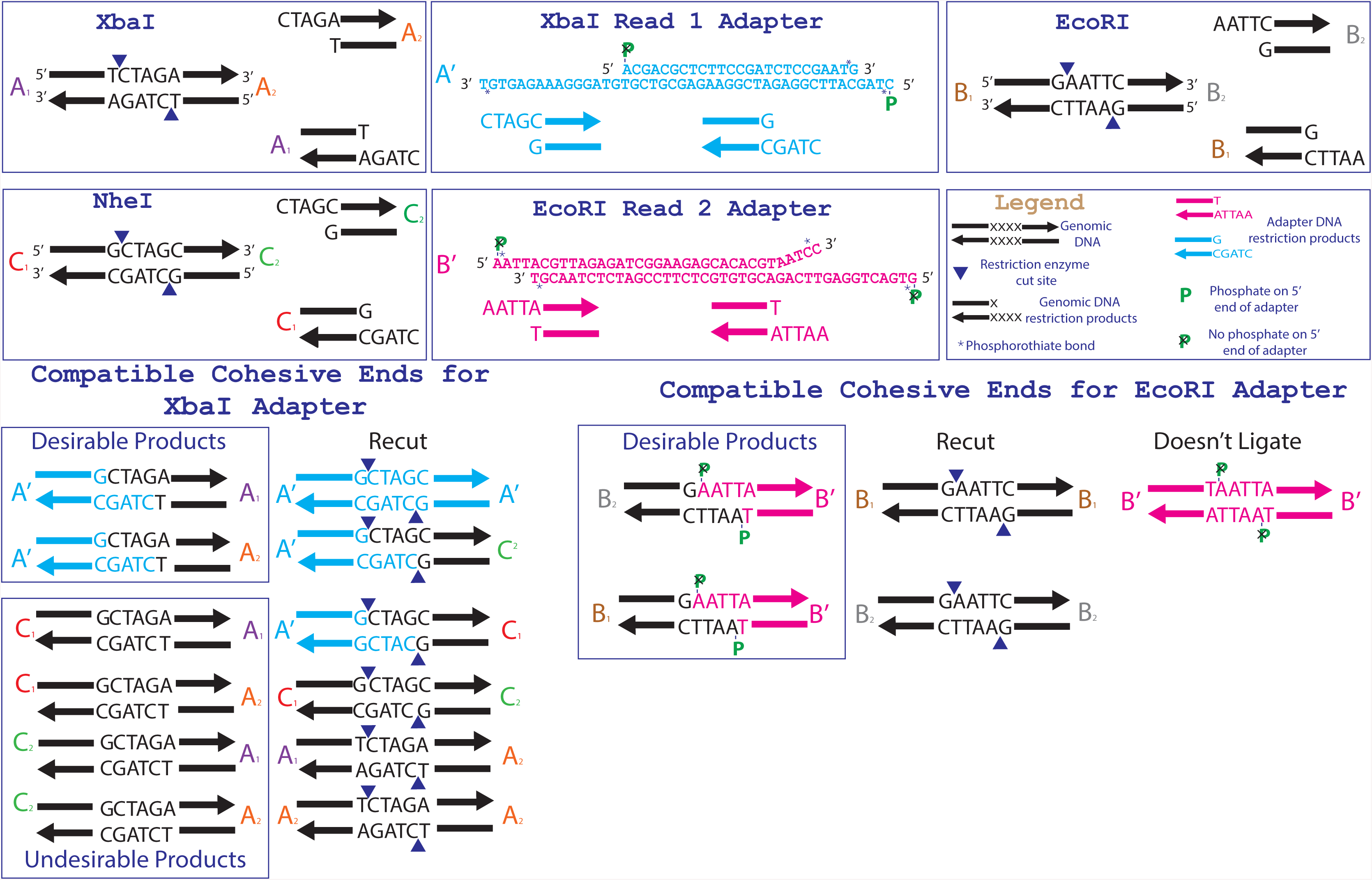
Specific adapter sequences and products created during the ligation of 3RAD libraries. The full adapter sequences for the 3RAD enzyme combination NheI, XbaI and EcoRI-HF (Table 1) are given in the top center boxes. The relevant recognition sequences for the three restriction endonucleases are given in the top outer boxes. The products that are formed from ligation of the triple-enzyme digests and adapters are shown at the bottom.

As described above, we cycle temperatures in the simultaneous digestion and ligation to allow this third RE to cut apart adapter-dimers, which increases the consistency and efficiency of 3RAD library preparation, even with limited amounts of sample DNA. However, while genomic DNA cut by this third RE and ligated to Read 1 adapters should be recut by the same RE, any remaining molecules of this form are suitable for PCR amplification and may be present in final libraries. 2RAD functions without this third RE, and in practice, the differences between 2RAD and ddRAD are: a) simultaneous digestion and ligation reactions; b) inclusion of variable-length internal indexes on each end (see below); c) compatibility with iTru primers from *Adapterama I* that allow for highly-multiplexed libraries to be pooled for sequencing on Illumina platforms (File S3); and d) potential substitution of the normal iTru5 primer containing specific index for iTru5-8N primer pool with 65,536 indexes, which facilitates identification and removal of PCR duplicates (Hoffberg et al., 2016).

Additionally, we constructed 2RAD/3RAD adapters so that each double-stranded adapter has one active strand (i.e., the bottom strand as shown in all figures herein) and one unused strand (i.e., the top strand in all figures herein). The dummy strand is simply used for structural support and correct 3D structure of the adapters and constructs through the ligation process. Both strands fit together on each side of the input DNA during ligation, but the nick between the sample DNA and Read 2 adapter top dummy strand is not ligated (Fig. S1). Thus, the unused strand construct breaks apart during PCR steps, and only those constructs with bottom strands that successfully ligate both kinds of adapters are amplified. This ensures that valid constructs with the correct restriction sites at opposite ends dominate the amplified library pools.

Additionally, the oligonucleotides for the top strand (as depicted in Figs. 1, S1, and S2) are not full-length, so they cannot be used as templates for the iTru5 or iTru7 primers. Finally, the top strand of the Read 2 adapter ends in five non-complementary bases so that it cannot serve as an unwanted primer during library amplification.

We achieved our next three design goals (i.e., 5–7) by including variable-length internal indexes—also known as “in-line barcodes” (Andrews et al., 2016)—within the Read 1 and Read 2 adapter stubs and making the adapter stubs compatible with the primers of Glenn et al. (2016; Figs. 1 and S1). For each adapter stub design, we have made eight versions of the Read 1 adapter stub and 12 versions of the Read 2 adapter stub (File S3). Each adapter stub version includes an internal index of 5, 6, 7, or 8 nucleotides (nt). The purpose of these internal indexes is twofold: 1) combinations of the Read 1 and Read 2 adapters create 96 (8 × 12) index combinations, which facilitates pooling of samples from 96-well plates (File S3); and 2) the variable length of each index increases base diversity within pools of libraries (Krueger, Andrews, & Osborne, 2011), which is important when sequencing libraries derived from RE digestion (Mitra et al., 2015; Glenn et al., 2016).

After ligating adapters, we create full-length libraries through reduced cycle PCR using the iTru5 and iTru7 primers of Glenn et al. (2016; Figs, 1, 3, S1, and S2). Because the 2RAD/3RAD adapters already include internal tags that can identify all samples in a 96-well plate, samples can be pooled prior to PCR and externally tagged with the iTru5 and iTru7 primers (to identify multiple plates of samples), or users can PCR amplify individual wells with unique external tags and pool PCR products, creating redundant indexing (Fig. 3). The resulting libraries are then size-selected and sequenced using the four standard Illumina TruSeq primers, each of which returns a different indexing read (Fig. 4). Because 2RAD/3RAD libraries are constructed using the iTru primers from *Adapterama I* (Glenn et al., 2016), they are compatible with (i.e., can be pooled with) iTru and Illumina TruSeq libraries prior to sequencing on Illumina sequencing instruments, achieving our final design goal. This potential for highly-multiplexed pools is especially important with the advent of platforms such as the Illumina NovaSeq, which is capable of producing up to 3000 Gb of data from one flow cell.

**Figure 3.**
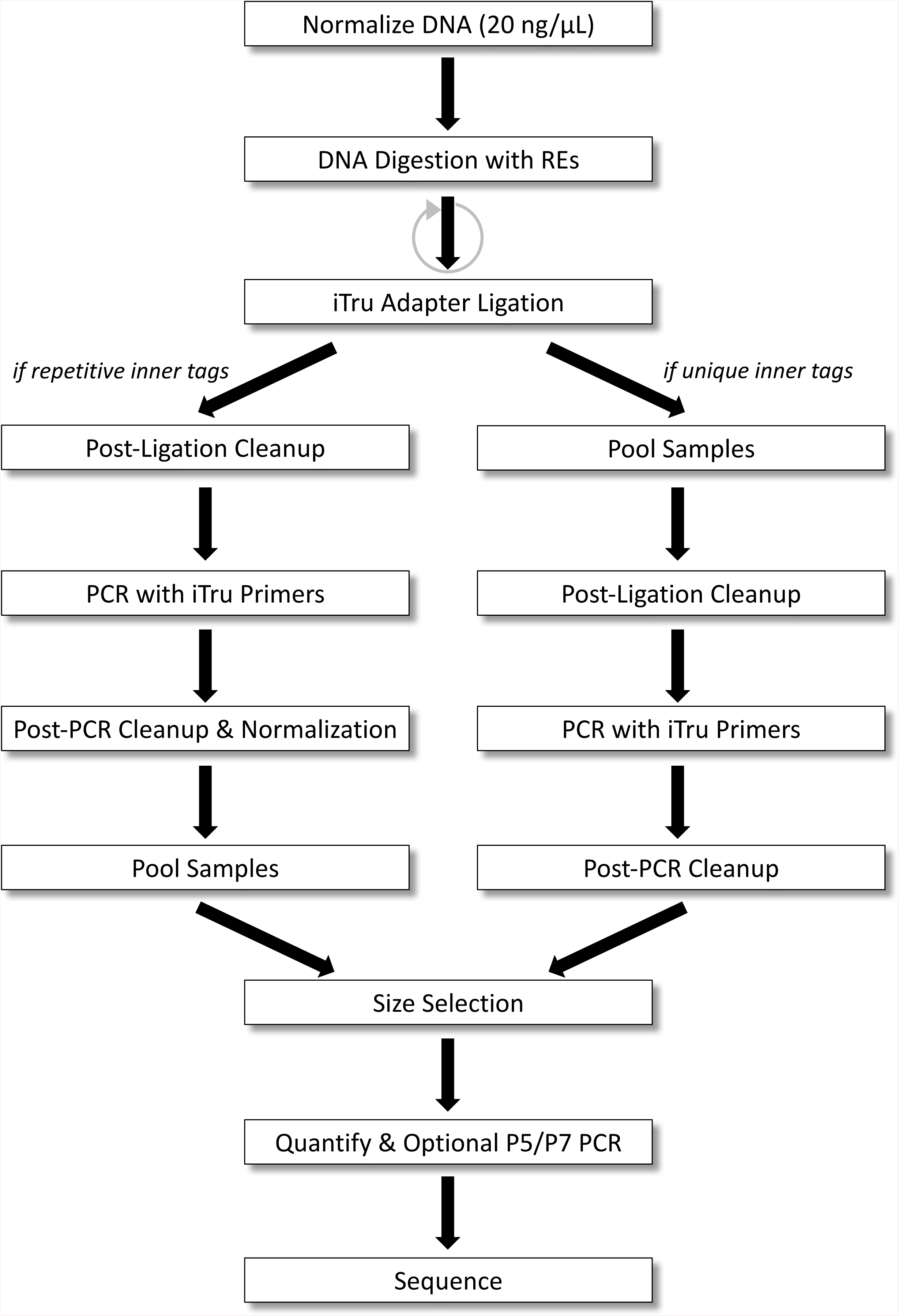
3RAD workflow for samples with unique or repeated adapter tags. DNA is normalized, digested with REs, and ligated to adapters. If indexes within adapters uniquely identify all samples (right), samples can be pooled before clean-up and PCR. If indexes do not uniquely identify individuals, PCR must be done separately on each sample, and samples must be normalized and cleaned before pooling. Then, samples are size-selected and quantified to determine if a final P5/P7 PCR should be performed before sequencing.

**Figure 4.**
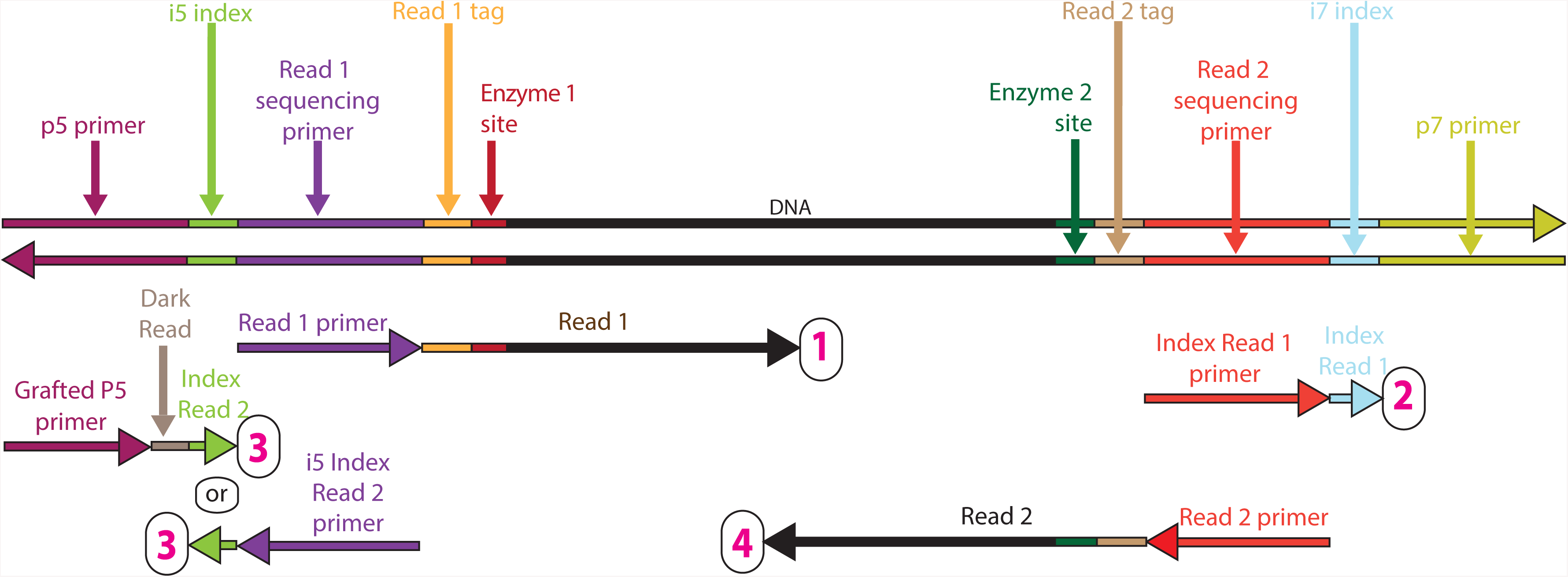
Sequencing reads that can be obtained from full length 3RAD library molecules. The top double stranded molecule shows a 3RAD library molecule prepared as described in the text (File S1). The horizontal arrows beneath the library molecule indicate Illumina sequencing primers (binding to the complementary strand of the library molecules). The tip of the arrowhead indicates the 3’ end of the primer and the direction of elongation for sequencing. Four sequencing reads are shown for each library prepared molecule, with one read for each index and each strand of the genomic DNA, including internal indexes. Reads are arranged 1 to 4 (numbered in magenta) from top to bottom, respectively. The arrow immediately 3’ of the primers, indicates the data that are obtained from that primer, with coloring that is consistent with 3RAD library molecule.

### 3RAD efficiency

We tested and compared our 2RAD and 3RAD protocols with the traditional ddRAD protocol using the same REs, adapters, and primers. To simplify the comparison between protocols, we used the pUC19 vector, which contains XbaI cut-site at position 423 and EcoRI cut-site at position 396, as template DNA. First, we amplified an approximately 500 bp fragment within the vector using the primers pUC19-215F-AAGGAGAAAATACCGCATCAGG and PUC19-774R-TAACCGTATTACCGCCTTTGAG. Then, we made four 10-fold dilution series. We used these five products (one stock and four dilutions) plus a negative control as input for 2RAD, 3RAD, and ddRAD (i.e., 2RAD with sequential digestion and ligation) libraries (File S5).

### 3RAD applied case studies

We tested our 3RAD protocol on eight example projects focused on diverse taxa: Kinosternidae (turtles), Ixodidae (ticks), *Eurycea bislineata* species complex (salamanders), *Wisteria floribunda* x *Wisteria sinensis* hybrid population (plants), *Rhodnius pallescens* (insects), *Gambusia affinis* (freshwater fish), *Sphyrna tiburo* (sharks), and *Sphyrna lewini* (sharks). Each dataset consisted of 12 to 24 samples. These projects span a broad diversity of organisms (e.g., in taxonomic classification, population size, heterozygosity level, and genome size), motivating evolutionary questions, and associated methods (i.e., from population genetics to phylogenetics; Table 3). After preliminary examination of several RE combinations, we used the REs XbaI, EcoRI-HF, and NheI and adapter sets R1.A and R2.1 (Design 1 in File S3) for all projects.

**Table 3.**
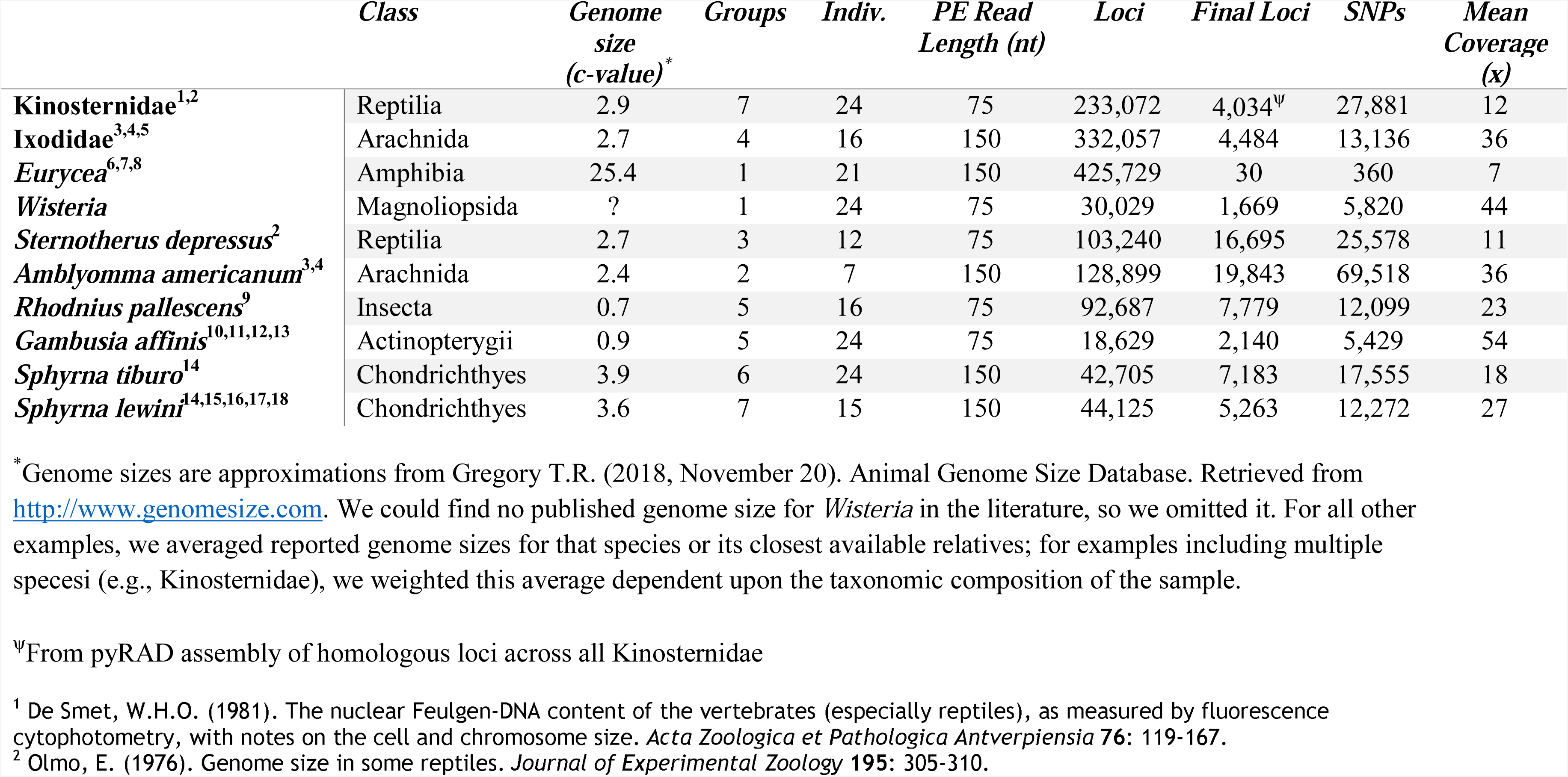
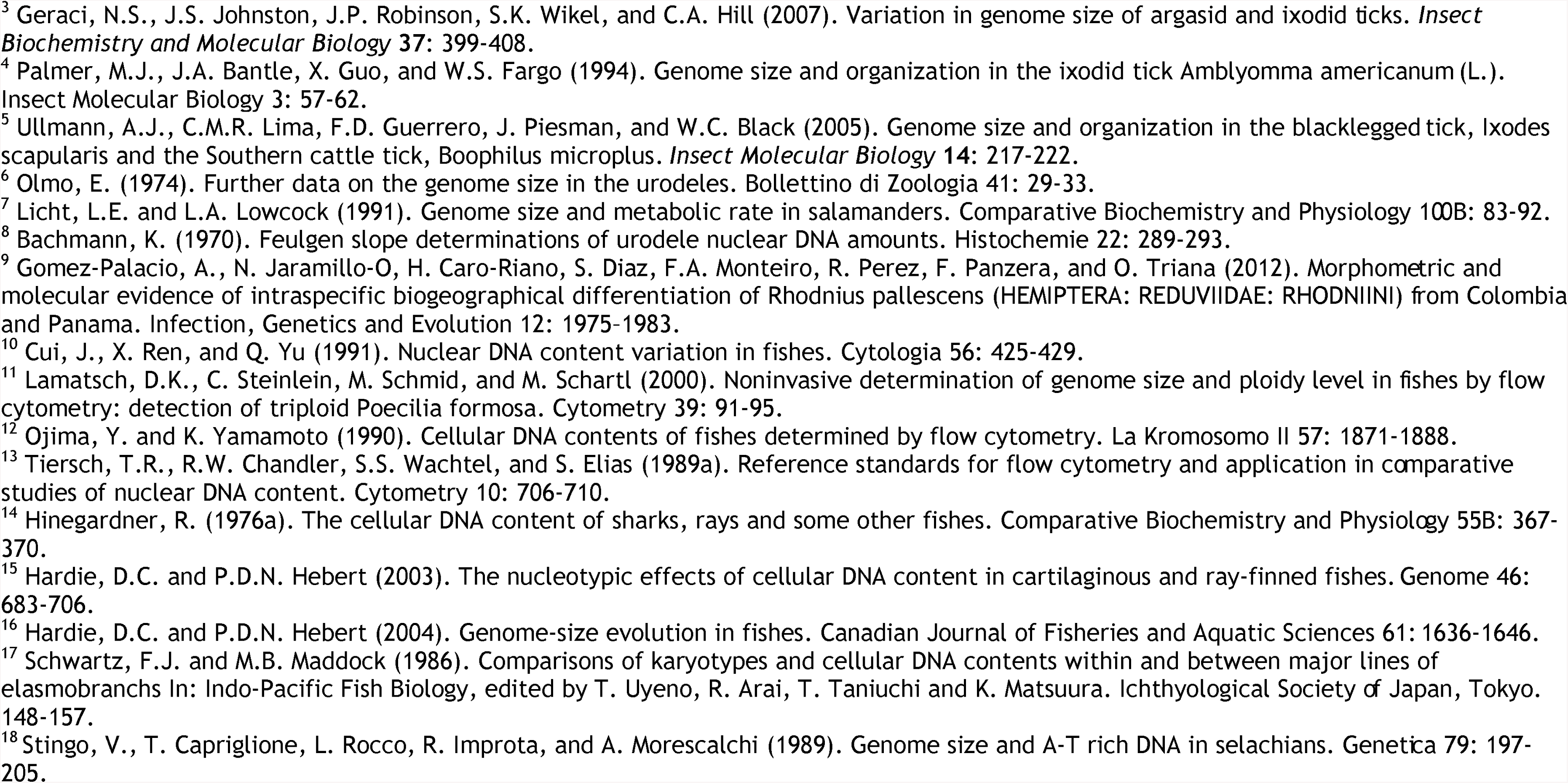
3RAD example projects. Classification and genome size of taxa, number of samples tested for each, Illumina read length (nt), number of loci obtained after the assembly method, number loci and SNPs obtained after filtering by only polymorphic loci shared in at least 75% of samples, and the average coverage among loci and individuals. The number of loci can be quite large and certainty of homology variable with distantly related samples, particularly if they have large genomes.

|  | *Class* | *Genome size (c-value)\** | *Groups* | *Indiv.* | *PE Read Length (nt)* | *Loci* | *Final Loci* | *SNPs* | *Mean Coverage (x)* |
| --- | --- | --- | --- | --- | --- | --- | --- | --- | --- |
| **Kinosternidae**<sup>1,2</sup> | Reptilia | 2.9 | 7 | 24 | 75 | 233,072 | 4,034<sup>ψ</sup> | 27,881 | 12 |
| **Ixodidae**<sup>3,4,5</sup> | Arachnida | 2.7 | 4 | 16 | 150 | 332,057 | 4,484 | 13,136 | 36 |
| **Eurycea**<sup>6,7,8</sup> | Amphibia | 25.4 | 1 | 21 | 150 | 425,729 | 30 | 360 | 7 |
| **Wisteria** | Magnoliopsida | ? | 1 | 24 | 75 | 30,029 | 1,669 | 5,820 | 44 |
| **Sternotherus depressus**<sup>2</sup> | Reptilia | 2.7 | 3 | 12 | 75 | 103,240 | 16,695 | 25,578 | 11 |
| **Amblyomma americanum**<sup>3,4</sup> | Arachnida | 2.4 | 2 | 7 | 150 | 128,899 | 19,843 | 69,518 | 36 |
| **Rhodnius pallescens**<sup>9</sup> | Insecta | 0.7 | 5 | 16 | 75 | 92,687 | 7,779 | 12,099 | 23 |
| **Gambusia affinis**<sup>10,11,12,13</sup> | Actinopterygii | 0.9 | 5 | 24 | 75 | 18,629 | 2,140 | 5,429 | 54 |
| **Sphyrna tiburo**<sup>14</sup> | Chondrichthyes | 3.9 | 6 | 24 | 150 | 42,705 | 7,183 | 17,555 | 18 |
| **Sphyrna lewini**<sup>14,15,16,17,18</sup> | Chondrichthyes | 3.6 | 7 | 15 | 150 | 44,125 | 5,263 | 12,272 | 27 |

\*Genome sizes are approximations from Gregory T.R. (2018, November 20). Animal Genome Size Database. Retrieved from http://www.genomesize.com. We could find no published genome size for *Wisteria* in the literature, so we omitted it. For all other examples, we averaged reported genome sizes for that species or its closest available relatives; for examples including multiple species (e.g., Kinosternidae), we weighted this average dependent upon the taxonomic composition of the sample.
<sup>ψ</sup>From pyRAD assembly of homologous loci across all Kinosternidae
<sup>1</sup> De Smet, W.H.O. (1981). The nuclear Feulgen-DNA content of the vertebrates (especially reptiles), as measured by fluorescence cytophotometry, with notes on the cell and chromosome size. *Acta Zoologica et Pathologica Antverpiensia* **76**: 119-167.
<sup>2</sup> Olmo, E. (1976). Genome size in some reptiles. *Journal of Experimental Zoology* **195**: 305-310.

### Library preparation

We prepared all libraries using similar methods, but some details varied among projects (Files S1 and S7). We used a modification of our method to incorporate molecular ID tags for the *Wisteria* project (see Hoffberg et al., 2016; Files S6 and S7), but we otherwise used the 3RAD library preparation method described above (Files S1, S2, and S7). Briefly, we digested sample DNA with REs, ligated adapters and simultaneously digested dimers and chimeras with two alternating cycles of ligation-digestion, pooled those libraries that had unique internal indexes (only in *Wisteria, Gambusia* and Kinosternidae projects), and purified ligation products. To generate full-length libraries, we performed a PCR using iTru primers that contained unique indexes (i.e., external indexes) to further differentiate individual samples (i.e., *Sphyrna*, Ticks, *Eurycea*, and *Rhodnius pallescens*) or projects (i.e., *Wisteria, Gambusia* and Kinosternidae), and purified libraries (Fig. 1). We quantified, pooled libraries, and size-selected libraries to capture fragments at 550 bp +/-10%. We quantified the resulting libraries, and in some cases, performed a limited-cycle PCR with P5 and P7 primers to increase library concentration before sequencing.

### Sequencing and data analyses

We sequenced libraries on multiple Illumina platforms in multiple core labs. We used the Illumina NextSeq 500 platform to generate PE75 data for the *Rhodnius, Gambusia*, Kinosternidae, and *Wisteria* projects and Illumina HiSeq 2500 or NextSeq 500 platforms to generate PE150 data for the Ixodidae, *Sphyrna,* and *Eurycea* projects. We planned all sequencing runs to produce approximately 1 million reads per sample, which facilitates comparison of the 3RAD results among species with varying genome sizes, with the exception of the Ixodidae project, where samples averaged 4 million reads.

We assembled data from each project independently using Stacks v1.42 (Catchen et al., 2013; Catchen et al., 2011; see File S6). For the *Wisteria* project, we used molecular ID tags to facilitate PCR duplicate removal with the module *clone_filter* in Stacks (Catchen et al., 2013; Hoffberg et al., 2016). We describe detailed parameters and software specifications for each project in File S6. Briefly, for most projects, we used the *process_radtags* program to demultiplex and/or clean and trim the sequence data. We parallel-merged the mates of paired-end reads. We used the *denovo_map* program assemble reads *de novo* and to calculate coverage, number of loci, and number of SNPs recovered for each project; we compared these data to genome size and sequencing read length (PE75 or PE150). Finally, we used the *populations* program to export loci shared in at least in 60-75% of localities and individuals to VCF files. Because there exists a reference genome for *Gambusia affinis* (Hoffberg et al. 2018; NCBI NHOQ01000000; details in File S6), we also assembled data from this project against the reference. For population-level datasets, we calculated *F*-statistics and performed preliminary population clustering analyses in Structure v2.3.4 (Pritchard, Stephens, & Donelly, 2000; File S6). For the Kinosternidae project, we conducted a *de novo* locus assembly using pyRAD v1.0.4 (Eaton, 2014; details in File S6).

Finally, we estimated the prevalence and impact of loci with third RE cut-sites in our data. We estimated the proportion of these third RE cut-sites relative to the first RE cut-site (i.e., intended cut-site) for five of the projects. To evaluate the effect of these loci in downstream analyses, we reanalyzed data from two of our projects (i.e., both *Sphyrna*) after removing third RE loci from the datasets. To do this, we reassembled data in Stacks v1.44 (Catchen et al., 2011; Catchen et al., 2013) using *process_radtags* two independent times: first, “rescuing barcodes”, cleaning, and trimming the raw sequence data as before, but disabling rad check (--disable_rad_check) to leave the cut-sites intact; and second, using the previous step’s output as input, checking only for exact, intended RE cut-sites (i.e., XbaI and EcoRI). From this output, we assembled and analyzed data similar to above, as detailed in File S6.

## Results

We developed four sets of adapters, each with eight versions of Read 1 adapters and 12 versions of Read 2 adapters (File S3). We modified the iTru_R2_5 index sequence for BamHI because this index creates a BamHI recognition site in the adapter; otherwise, all adapters use a universal set of index sequences. The cost of synthesizing oligonucleotides for the adapters varies with synthesis scale, but it starts as low as ∼$350 (US) per design set, with a recommended scale (100 nmol) costing ∼$500 (US) per design set when synthesized into 96-well plates, which are sufficient for up to ∼4800 sample libraries.

3RAD libraries can be constructed routinely with approximately 12 hours of hands-on time over the course of 2–3 days, with some variation depending mostly upon the step at which samples are pooled. The initial cost of restriction digestion and ligation is ∼$0.85 per sample (File S3, “Library_prep_costs” Sheet). If PCRs are conducted individually, this adds > $1 per sample, but if the ligations are pooled before PCR, then this cost averages to $0.06 per sample. Size-selection using the Pippin Prep adds $0.12 per sample, assuming user access to the equipment. A total of $0.25 per sample is required for tips, plates and tubes. Thus, when samples are pooled prior to PCR, the total cost for library preparation is about $1.35 per sample.

We demonstrate that 3RAD libraries have fewer adapter-dimers and a higher concentration of library constructs than 2RAD or ddRAD libraries, which is particularly important when libraries are constructed with low input DNA concentrations (Fig. 5; File S5). This improved performance is a consequence of maximizing the efficiency of the ligation of the adapters to library fragments.

**Figure 5.**
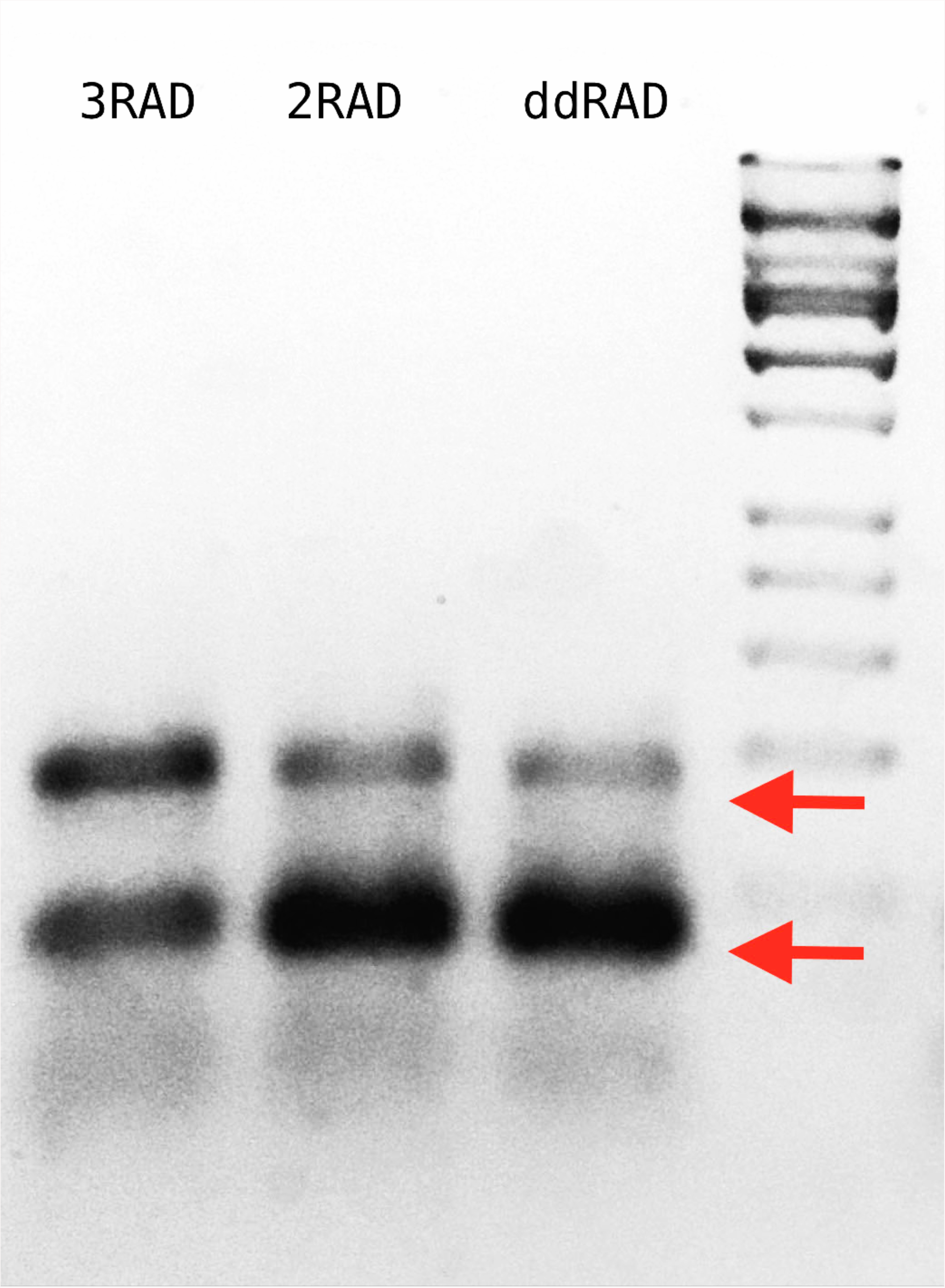
Agarose gel with 3RAD, 2RAD, and ddRAD library products performed on pUC19 vector with an input quantity of 0.05 ng. The band close to the 200 bp size standard (arrow above) is that corresponding to a proper library construct. The band below the 100 pb size standard (arrow below) corresponds to adapter-dimers (File S5). The gel indicates that 3RAD libraries outperformed the other two types of libraries tested by decreasing the adapter-dimers and therefore increasing the quantity of desired library constructs.

In the Ixodidae example project, we obtained between 1.8–7.3 (mean = 4.2) million reads per sample, and for all other projects, we obtained between 0.6–3.6 (mean = 1.3) million reads per sample. With the exception of one sample from the Ixodidae project, we always recovered a high percentage of retained reads (78.9–99.7%) after cleaning and filtering steps (Table S1). Average coverage per locus varied from 6x for *Eurycea* to 70x for *Gambusia* (Table 3; Table S1).

Our initial assemblies contained between 18,629 loci (*Gambusia*) and 425,729 loci (*Eurycea*). After filtering to retain only polymorphic loci found in at least 75% of individuals within each population, we recovered between 30 loci (*Eurycea*) and 19,843 loci (Ixodidae) containing between 360 and 69,518 SNPs, respectively (Table 3). As expected for RADseq protocols, the number of loci we obtained in the initial steps was proportional to the genome size, with more loci recovered in organisms with larger genomes. Due to the manner in which we filtered these loci, the final number of loci recovered is dependent upon the intrinsic genetic variability of the organism, the scale of sampling, and sequencing coverage (Fig. S3). Detailed results for each project can be found in File S6.

Third RE loci were present in all datasets, comprising an average of 20.5% (sd = 14.1%) of all reads. The percentage of reads from third RE loci varied both among and within datasets, showing a nonrandom pattern with respect to R1 adapter index used (Fig. S4). Removing third RE loci from our raw reads increased mean coverage of remaining loci (Tables S2 and S3) and reduced the size of our final datasets from 7,183 to 6,738 loci in *Sphyrna tiburo* and 5,263 to 4,807 loci in *Sphyrna lewini.* Estimates of *F*_ST_ and results from Structure were qualitatively similar in analyses including and excluding third RE loci (Table S4).

## Discussion

We present an efficient, flexible, and low-cost system for preparing dual-digest RADseq. Our method uses the iTru primers from *Adapterama I* (Glenn et al., 2016) and modifies the adapter stub for RADseq by adding the appropriate overhang bases, as well as variable-length internal indexes which facilitates a single Illumina lane to be shared by many quadruple-indexed libraries. To illustrate the utility of our method, we present summary statistics from analyses of eight small example projects, representing diversity in taxonomy and scientific objectives. For each project, we obtained thousands of loci containing SNPs for downstream population genetic and phylogenetic analyses. Among the example projects, we highlighted the role of genome size, genetic variation, and sequencing coverage in determining the quality and quantity of data recovered. For example, when processed differently, we recovered large numbers of homologous loci both among species in the family Kinosternidae and within a single representative species, *Sternotherus depressus,* from the same libraries and sequencing reads. These data are informative both for studying variation among populations of *S. depressus* and across relatively deep evolutionary time (∼55 Myr; File S6; Scott, Glenn, & Rissler, 2017). We note that when datasets span deeper evolutionary time (e.g., *Eurycea* and Kinosternidae), we recover more loci in initial steps, but fewer of these loci are shared among individuals. However, many studies support the utility of these large and sparse data matrices for phylogenetic studies (e.g., Streicher, Schulte, & Wiens, 2015; Hosner et al., 2016). For detailed discussion of the analyses for each dataset, see File S6.

By using the same set of REs on DNA from a diverse set of organisms, we have demonstrated that our 3RAD method recovers suitable numbers of loci and SNPs from organisms with varying genome characteristics. As expected, when we use the same set of REs (and thus similar expected frequency of cut-sites) and the same size-selection criteria for organisms with varying genome size, the average sequencing coverage per locus decreases as genome size increases (Fig. 6). For example, our dataset generated here from *Eurycea* (C-value = ∼25; Table 3) produced few loci meeting our coverage criteria, but higher sequencing coverage can remedy this. Alternatively, the number of loci in libraries can be tuned by changing REs or size-selection criteria (Peterson et al., 2012). Our initial testing of all four design sets from the 3RAD method on chicken DNA showed that the number of loci varied as expected (i.e., REs with shorter recognition sequences yielded more loci; data not shown). Additionally, using broad size-distributions in the size-selection step retains more loci and allows for greater size tolerance among alleles, whereas narrow distributions yield fewer loci. We suspect that use of narrow size-distributions excludes alleles at loci with significant size variation and may lead to increased levels of incorrect genotype calls due to the missing alleles outside of the selected size-range, but because most researchers are targeting loci without size variation, this bias should be small. A more significant issue related to narrow size-distributions is that a significant proportion of loci will be near the size cut-off and coverage decreases for these loci because some molecules of the targeted size will not be among the fragments retained (e.g., simply due to variance in migration through a gel).

**Figure 6.**
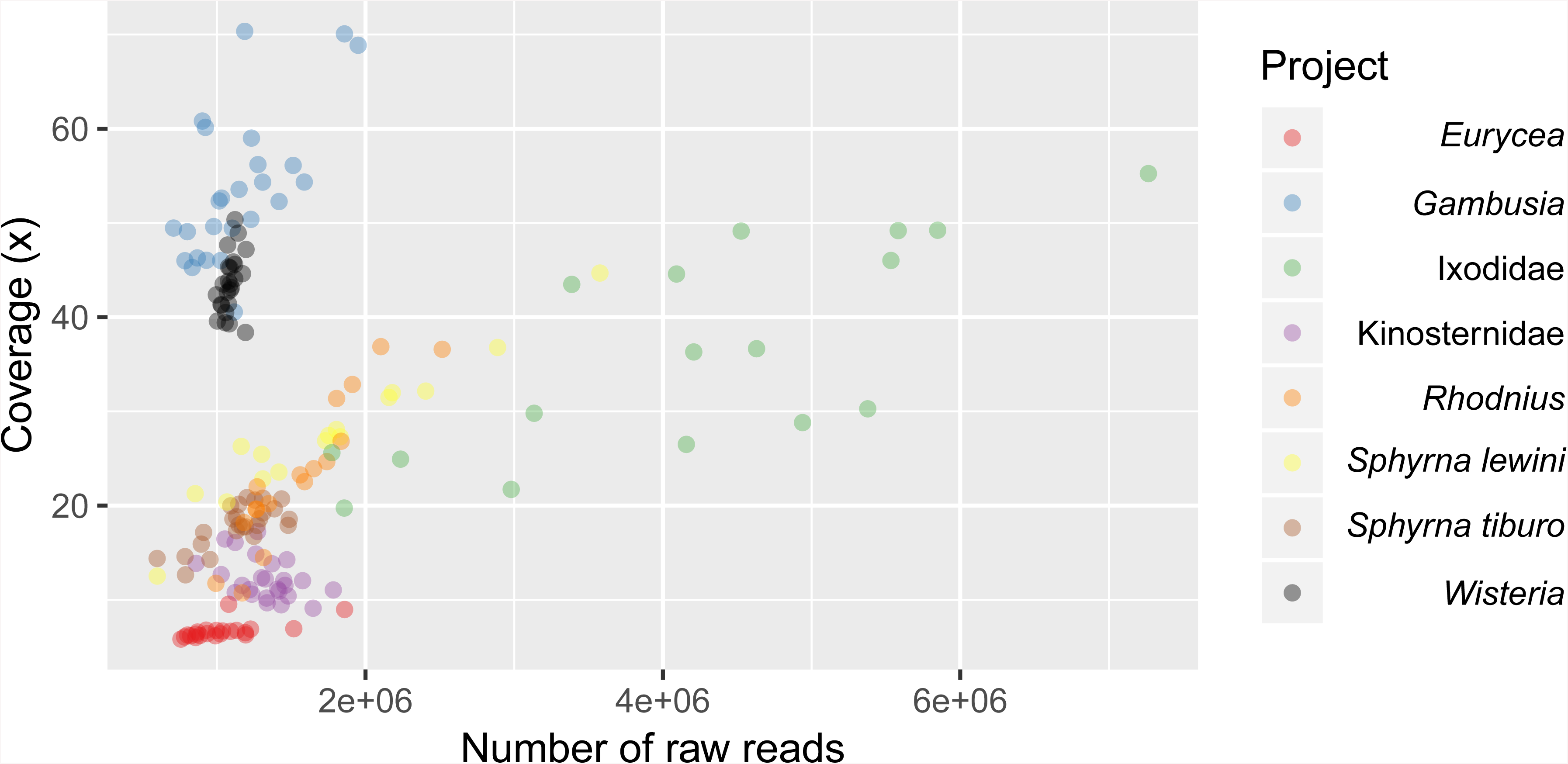
Scatterplot of the average coverage of all loci (polymorphic and fixed) for each sample relative to sequencing depth of each sample. *Eurycea* have the largest genome size and therefore the lowest average coverage per locus with approximately 1,000,000 reads. Average coverage increases as the genome size decreases (Fig. S3).

A key advance of the 3RAD workflow (Files S1-S2) is the combination of enzymes and adapters used during digestion, ligation, and PCR steps to create the desired construct while minimizing the presence of dimers, chimeras, and improperly formed library molecules (those lacking restriction sites at both ends). Sequential and simultaneous digestion and ligation without buffer exchange increases the efficiency of the lab workflow and decreases the amount of input DNA required. The 3RAD adapters we designed function with multiple REs that leave compatible, cohesive ends (Table 2). Thus, there are at least 72 different RE combinations possible with the current adapter sets. Although this flexibility is desirable, having adapters compatible with multiple REs means that it was necessary to name the adapters based on the third RE, which can be confusing because the third RE is *not* the desired restriction site in the sample libraries (and their resulting reads). The design spreadsheet (File S3, Sheets: Design_1-4) can be easily modified to accommodate other REs to create additional designs. These sheets also incorporate cost calculations so that researchers may easily change the costs to reflect updated pricing from their supplier(s).

It should be noted that certain RE combinations work well with some species, but not with others, primarily due to the presence of restriction sites within repetitive elements. Thus, our standard strategy is to empirically determine what RE combination is best for any particular organism in a few representative samples. We start by looking at the distribution of post-PCR library DNA run through an agarose gel and exclude RE combinations that don’t produce even smears or that have dense bands in the desired size range. Sometimes, we then size-select and sequence libraries from one or two RE combinations from this small batch of samples to determine which combination produces the most variable loci.

Our results show that XbaI, EcoRI-HF and NheI provide a suitable combination of REs for a wide range of organisms, including plants, vertebrates, and invertebrates. We have used all of the adapter designs and many other RE combinations from the enzyme list (Table 1) to survey SNPs in a variety of organisms. While not all combinations work well in all organisms, most of the organisms we have studied to date work well with REs from Design 1 or Design 2 adapters. Although viable, we only rarely use Designs 3 and 4 (R1.C, R1.D, R2.3, and R2.4).

In our standard protocols (Files S1 and S5), we digest DNA with two different REs (2RAD) or three different REs (3RAD) to create sticky ends for adapter ligation (similar to ddRAD and 2-enzyme GBS). In 3RAD, the third RE digests a recognition site formed by self-ligation of the phosphorylated adapters (Fig. 2). Although the third RE facilitates creation of libraries with very little input DNA (≤ 0.1 ng; File S5), it does come with a cost. The third RE also cuts genomic DNA that can be ligated to the R1 adapter, but the adapter:DNA ligation product is susceptible to re-cleavage by the RE. To encourage this, our digestion/ligation cycling ends with a digestion step to cleave as many of these products as possible. Still, an average of 20% of loci in our assemblies had these third RE cut-sites, and the nonrandom relationship between R1 adapter index and the prevalence of these loci suggests that the index has an effect upon the efficiency of the third RE in cleaving the re-created recognition site. These loci are, in principle, suitable for downstream analyses, but because the protocol is designed to minimize their retention, they should have lower coverage than those with the intended RE cut-site. A high prevalence of these off-target loci can require additional sequencing (and thus, increase costs), but these reads can easily be filtered and removed for all downstream analyses if desired. The third RE is not required for this procedure and further investigation into to trade-offs of including this RE (i.e., 2RAD vs. 3RAD) and its optimal concentration are warranted. Alternatively, it is possible to engineer adapters that can ligate to genomic DNA, but not self-ligate (e.g., using 3’ dideoxycytidine), which we have done. Unfortunately, the adapters are significantly more expensive and were not stable when stored for ≥ ∼6 months, both of which make the method impractical. Further research into other modifications or storage solutions for these adapters is warranted.

As detailed in Hoffberg et al. (2016), our 3RAD approach is easy to modify to include tags that allow the removal of PCR duplicates (File S7) through a process comparable to quaddRAD (Franchini et al., 2017). Our molecular ID tag protocol, which uses an iTru5-8N primer, can be used to make libraries for different types of downstream processes and/or modified for other types of libraries. For example, Hoffberg et al. (2016) used the iTru5-8N primer and sequence capture to focus sequencing on informative loci, reduce missing data, and remove PCR duplicates.

Our 2RAD/3RAD methods are similar to other dual-digest RADseq methods (e.g., ddRAD and 2-enzyme GBS), and most of the advantages of our general approach have been described previously (Andrews et al., 2016; Harvey et al., 2016; Heyduk et al., 2016). Our 2RAD/3RAD method achieves the seven methodological design objectives. We have demonstrated that the third RE increases ligation efficiency by reducing adapter-dimers; thus, much less input DNA is necessary. In addition, multiple REs are compatible with many of the adapters (Table 1), all indexes used herein conform a minimum edit distance of 3 (Faircloth & Glenn, 2012), we use limited PCR-cycles of pooled ligations to reduce PCR bias, and our method facilitates the easy incorporation of molecular ID tags to detect PCR duplicates in downstream analyses (Hoffberg et al., 2016).

## Conclusions

In summary, 2RAD/3RAD protocols function similar to other dual-digest RADseq methods but are easier to perform, have lower startup costs, have increased utility with low-concentration input DNA, and produce libraries that can be highly-multiplexed and pooled with other Illumina libraries.

## Acknowledgements

Portions of the oligonucleotide sequences are © 2007-2019 Illumina, Inc., all rights reserved. Derivative works, such as the oligonucleotides presented here, are authorized for use with Illumina instruments and products only. All other uses are strictly prohibited. We thank Julian Catchen for modifying Stacks and Isaac Overcast and Deren Eaton for modifying pyRAD to deal with the variable-length dual internal tags, as well as for incorporating additional enzymes to facilitate data processing of 3RAD data. We thank Erin Lipp for generously sharing laboratory space and equipment. We thank Jane Stewart, Marin Talbot Brewer and Bryan Carstens for suffering through the primal 2RAD development with modified bases in the adapters, our colleagues at the Georgia Genomics and Bioinformatics Core for conducting Illumina sequencing, the Georgia Advanced Computing Resource Center and our colleagues there for support, and Chad Locklear of Integrated DNA Technologies for his assistance with and support for adapter oligonucleotide development and synthesis. We also thank our colleagues involved in the logistics and sample collection for example projects included in this paper.

---

Supplementary Figure S1

**Specific reactions and sequence constructs created by the 2RAD/3RAD library workflow** Detailed sequences for the workflow displayed in Fig. 1. 5’ phosphates are indicated with a red “P”. The Read 2 adapter lacks phosphates; thus, when it is ligated to the digested genomic DNA, a nick remains in the top strand (i.e., the phosphodiester bond between the genomic DNA and the adapter is missing, indicated by the “P” with an “X” through it).

Supplementary Figure S2

**Products created during the first three rounds of PCR during 2RAD/3RAD library construction**

This figure demonstrates how only the bottom strand is used to form the fully-functional 3RAD libraries.

Supplementary Figure S3

**Scatterplot of the normalized coverage against the genome size for sample project**

We divided the average coverage for each sample by the total number of retained reads for that sample to obtain the normalized coverage. The *Wisteria* dataset is not included because the genome size is unknown.

Supplementary Figure S4

**Boxplot of the proportion of reads with NheI cut-sites in eight of the sample projects**

The proportion of the total raw R1 reads derived from loci cut by NheI, rather than XbaI, in eight of our sample projects broken down by which adapter (i.e., NheI_A-H) was used.

Supplementary File S1

**High-Throughput 3RAD Protocol**

Step by step library construction for 3RAD libraries considering both, samples pooled after ligation because all adapter combinations are used and, samples that are pooled after PCR.

Supplementary File S2

**3RAD video presentation, what is happening inside the tube?**

This presentation demonstrates the key features of 3RAD, including how the adapters perform during ligation and PCR, how the adapters and primers can be used combinatorically, and both the desired and undesired ligation products that can be formed during 3RAD. This presentation is also available at: https://www.youtube.com/watch?v=ZOmwOtfP3N4

Supplementary File S3

**3RAD iTru adapter designs, indexed iTru primers, and detailed library prep costs per sample**

Excel workbook containing sheets for each of the four pairs of adapter designs. The designs are paired within the workbook to save space, but each design is independent, thus design 1 Read 1 adapters can be used with design 2 Read 2 adapters. Each adapter requires two oligonucleotides to form a partially complementary pair. Also, we present an example of the iTru primers with incorporated indexes to use in 3RAD libraries. And finally, tab with costs per sample for library prep, considering each step in the process.

Supplementary File S4

**How to handle plates with 3RAD adapter aliquots**

Document with instructions of how to reconstitute and anneal 3RAD dried adapters to further use in 3RAD libraries.

Supplementary File S5

**3RAD efficiency compared to ddRAD**

Step by step protocol with instructions on how the comparison of 3RAD, 2RAD and ddRAD methods was surveyed on pUC19 vector.

Supplementary File S6

**Supplementary methods and results for 3RAD example datasets**

A detailed guide through the methods, results, and discussion of sequence analyses from 3RAD data generated for each example dataset presented in this manuscript.

Supplementary File S7

**3RAD Libraries with Molecular ID Tags Protocol**

Step by step protocol with instructions to build 3RAD libraries with molecular ID tags to detect PCR duplicates, using iTru5 8N primer (See Hoffberg *et al*., 2016)

The authors declare competing interests. The EHS DNA lab provide oligonucleotide aliquots and library preparation services at cost, including some oligonucleotides and services that make use of the adapters and primers presented in this manuscript (baddna.uga.edu). The information we present allows all researchers to synthesize the oligonucleotides at any vendor of their choice, follow or modify the library preparation techniques we have included, and freely publish results simply with proper attribution of this paper and Illumina^®^™.

This work was partially supported by DEB-1242241, DEB-1242260, DEB-1136626, DEB-1146440, DEB-1353815, DEB-1405599, DGE-0903734, DGE-1452154, and OISE 0730218 from the U.S. National Science Foundation. Data collection for the hammerhead shark was supported by the DGAPA (PAPIIT IG201215).

