## Supplementary material for "Adapterama III: Quadruple-indexed, double/triple-enzyme RADseq libraries (2RAD/3RAD)": Table S1

**Total number of reads per sample, reads retained after cleaning and filtering in Stacks, percent of reads retained, and average coverage over all loci**

Bold values correspond to averages for each project.

|  | Raw Reads | Retained Reads | % Retained Reads | Coverage (x) |
| --- | --- | --- | --- | --- |
| ***Gambusia affinis*** | **1149076** | **1137483** | **99.0** | **53.57** |
| Ga_af_01 | 1014974 | 1010844 | 99.6 | 52.34 |
| Ga_af_02 | 1031166 | 1026590 | 99.6 | 52.64 |
| Ga_af_03 | 784948 | 781906 | 99.6 | 45.99 |
| Ga_af_04 | 1232248 | 1227979 | 99.7 | 59.01 |
| Ga_af_05 | 922482 | 915340 | 99.2 | 60.14 |
| Ga_af_06 | 1857706 | 1842636 | 99.2 | 70.07 |
| Ga_af_07 | 800956 | 787397 | 98.3 | 49.08 |
| Ga_af_08 | 929254 | 916058 | 98.6 | 46.04 |
| Ga_af_09 | 1307694 | 1279267 | 97.8 | 54.34 |
| Ga_af_10 | 868072 | 856521 | 98.7 | 46.29 |
| Ga_af_11 | 1228392 | 1206904 | 98.3 | 50.38 |
| Ga_af_12 | 833852 | 821814 | 98.6 | 45.27 |
| Ga_af_13 | 1276016 | 1254565 | 98.3 | 56.20 |
| Ga_af_14 | 1418250 | 1380902 | 97.4 | 52.28 |
| Ga_af_15 | 902190 | 896034 | 99.3 | 60.82 |
| Ga_af_16 | 1949978 | 1935951 | 99.3 | 68.86 |
| Ga_af_17 | 1186866 | 1178099 | 99.3 | 70.34 |
| Ga_af_18 | 1512898 | 1496099 | 98.9 | 56.11 |
| Ga_af_19 | 1116982 | 1113619 | 99.7 | 40.55 |
| Ga_af_20 | 978550 | 975575 | 99.7 | 49.62 |
| Ga_af_21 | 1025566 | 1022213 | 99.7 | 46.01 |
| Ga_af_22 | 708264 | 705874 | 99.7 | 49.46 |
| Ga_af_23 | 1102860 | 1085145 | 98.4 | 49.45 |
| Ga_af_24 | 1587668 | 1582262 | 99.7 | 54.34 |
| **Kinosternidae** | **1325481** | **1294994** | **97.7** | **12.22** |
| St_tr_01 | 859958 | 855058 | 99.4 | 13.87 |
| St_tr_02 | 1122540 | 1116843 | 99.5 | 16.09 |
| Ki_ba_01 | 1053596 | 1048332 | 99.5 | 16.45 |
| Ki_ba_02 | 1261866 | 1246626 | 98.8 | 14.86 |
| St_mi_01 | 1273720 | 1218151 | 95.6 | 17.25 |
| St_mi_02 | 1337696 | 1302375 | 97.4 | 9.68 |
| St_pe_01 | 1169984 | 1142071 | 97.6 | 11.53 |
| St_pe_02 | 1219506 | 1207579 | 99.0 | 11.09 |
| St_od_01 | 1408406 | 1387826 | 98.5 | 11.13 |
| St_od_02 | 1457640 | 1423300 | 97.6 | 11.53 |
| St_ca_01 | 1449886 | 1431518 | 98.7 | 12.06 |
| St_ca_02 | 1028690 | 1003891 | 97.6 | 12.68 |
| St_de_01 | 1470502 | 1441759 | 98.0 | 14.25 |
| St_de_02 | 1782838 | 1732309 | 97.2 | 11.05 |
| St_de_03 | 1300368 | 1278841 | 98.3 | 12.34 |
| St_de_04 | 1478652 | 1450120 | 98.1 | 10.41 |
| St_de_05 | 1334344 | 1289642 | 96.6 | 10.19 |
| St_de_06 | 1235302 | 1167504 | 94.5 | 10.59 |
| St_de_07 | 1432174 | 1400775 | 97.8 | 9.47 |
| St_de_08 | 1417324 | 1378327 | 97.2 | 10.94 |
| St_de_09 | 1122922 | 1077956 | 96.0 | 10.77 |
| St_de_10 | 1647010 | 1621984 | 98.5 | 9.11 |
| St_de_11 | 1370474 | 1331399 | 97.1 | 13.83 |
| St_de_12 | 1576136 | 1525674 | 96.8 | 12.03 |
| ***Sphyrna tiburo*** | **1148868** | **1071967** | **93.3** | **17.95** |
| Sp_ti_01 | 1287858 | 1228641 | 95.4 | 18.60 |
| Sp_ti_02 | 1308450 | 1230170 | 94.0 | 20.79 |
| Sp_ti_03 | 1191718 | 1143276 | 95.9 | 17.75 |
| Sp_ti_04 | 1093198 | 1009118 | 92.3 | 19.98 |
| Sp_ti_05 | 1131494 | 1074561 | 95.0 | 17.34 |
| Sp_ti_06 | 1107376 | 1022300 | 92.3 | 18.58 |
| Sp_ti_07 | 1270442 | 1204857 | 94.8 | 17.90 |
| Sp_ti_08 | 1147090 | 1068668 | 93.2 | 20.14 |
| Sp_ti_09 | 1386368 | 1270284 | 91.6 | 19.66 |
| Sp_ti_10 | 894330 | 845769 | 94.6 | 15.91 |
| Sp_ti_11 | 1309184 | 1254979 | 95.9 | 19.24 |
| Sp_ti_12 | 1202502 | 1104610 | 91.9 | 20.85 |
| Sp_ti_13 | 1255266 | 1195822 | 95.3 | 20.59 |
| Sp_ti_14 | 1434898 | 1365345 | 95.2 | 20.71 |
| Sp_ti_15 | 1248378 | 1172229 | 93.9 | 16.74 |
| Sp_ti_16 | 910556 | 845323 | 92.8 | 17.16 |
| Sp_ti_17 | 788708 | 701804 | 89.0 | 12.65 |
| Sp_ti_18 | 953068 | 891180 | 93.5 | 14.28 |
| Sp_ti_19 | 1132048 | 1039236 | 91.8 | 18.81 |
| Sp_ti_20 | 1171024 | 1066234 | 91.1 | 17.73 |
| Sp_ti_21 | 785564 | 732048 | 93.2 | 14.59 |
| Sp_ti_22 | 597788 | 555368 | 92.9 | 14.39 |
| Sp_ti_23 | 1479190 | 1340062 | 90.6 | 17.91 |
| Sp_ti_24 | 1486342 | 1365324 | 91.9 | 18.55 |
| ***Sphyrna lewini*** | **1730896** | **1645522** | **95.1** | **26.88** |
| Sp_le_01 | 2404536 | 2324681 | 96.7 | 32.16 |
| Sp_le_02 | 2887858 | 2728601 | 94.5 | 36.78 |
| Sp_le_03 | 1829342 | 1763144 | 96.4 | 27.30 |
| Sp_le_04 | 2157066 | 2081759 | 96.5 | 31.49 |
| Sp_le_05 | 1301154 | 1026390 | 78.9 | 25.45 |
| Sp_le_06 | 3576314 | 3440441 | 96.2 | 44.69 |
| Sp_le_07 | 1804516 | 1706835 | 94.6 | 28.08 |
| Sp_le_08 | 1417000 | 1354153 | 95.6 | 23.56 |
| Sp_le_09 | 1308722 | 1251513 | 95.6 | 22.83 |
| Sp_le_10 | 1163354 | 1077005 | 92.6 | 26.29 |
| Sp_le_11 | 853748 | 807511 | 94.6 | 21.27 |
| Sp_le_12 | 2179644 | 2067373 | 94.8 | 31.99 |
| Sp_le_13 | 598658 | 571367 | 95.4 | 12.52 |
| Sp_le_14 | 1067186 | 1018289 | 95.4 | 20.39 |
| Sp_le_15 | 1730896 | 1645522 | 95.1 | 26.88 |
| ***Eurycea bislineata* complex** | **1038190** | **961101** | **92.6** | **6.69** |
| Eu_bi_01 | 1024050 | 963074 | 94.0 | 6.39 |
| Eu_bi_02 | 1191172 | 1127220 | 94.6 | 6.51 |
| Eu_bi_03 | 1517214 | 1380588 | 91.0 | 6.93 |
| Eu_bi_04 | 758056 | 648464 | 85.5 | 5.83 |
| Eu_bi_05 | 1193818 | 1139118 | 95.4 | 6.28 |
| Eu_bi_06 | 990288 | 921535 | 93.1 | 6.18 |
| Eu_bi_07 | 1226288 | 1139118 | 92.9 | 6.89 |
| Eu_bi_08 | 1858806 | 1606623 | 86.4 | 8.97 |
| Eu_bi_09 | 996126 | 965042 | 96.9 | 6.75 |
| Eu_bi_10 | 869316 | 825513 | 95.0 | 6.58 |
| Eu_bi_11 | 1091044 | 1018987 | 93.4 | 6.66 |
| Eu_bi_12 | 860568 | 748859 | 87.0 | 6.35 |
| Eu_bi_13 | 1132550 | 1099827 | 97.1 | 6.75 |
| Eu_bi_14 | 856816 | 821822 | 95.9 | 6.00 |
| Eu_bi_15 | 802696 | 744934 | 92.8 | 6.26 |
| Eu_bi_16 | 823006 | 721606 | 87.7 | 6.18 |
| Eu_bi_17 | 1079212 | 1020483 | 94.6 | 9.53 |
| Eu_bi_18 | 927524 | 883587 | 95.3 | 6.78 |
| Eu_bi_19 | 782614 | 733865 | 93.8 | 6.06 |
| Eu_bi_20 | 885804 | 774125 | 87.4 | 6.21 |
| Eu_bi_21 | 935024 | 898733 | 96.1 | 6.41 |
| ***Wisteria*** | **1088589** | **1074290** | **98.7** | **43.52** |
| Wi_sp_01 | 1042198 | 1033185 | 99.1 | 43.52 |
| Wi_sp_02 | 1082980 | 1059573 | 97.8 | 39.29 |
| Wi_sp_03 | 1110788 | 1100768 | 99.1 | 45.86 |
| Wi_sp_04 | 1072432 | 1064120 | 99.2 | 47.66 |
| Wi_sp_05 | 1070554 | 1063285 | 99.3 | 42.64 |
| Wi_sp_06 | 1090082 | 1080777 | 99.1 | 42.88 |
| Wi_sp_07 | 1055748 | 1042858 | 98.8 | 39.41 |
| Wi_sp_08 | 1119726 | 1107853 | 98.9 | 45.60 |
| Wi_sp_09 | 1031908 | 1014764 | 98.3 | 41.27 |
| Wi_sp_10 | 1027714 | 1004712 | 97.8 | 41.34 |
| Wi_sp_11 | 1003116 | 992424 | 98.9 | 39.59 |
| Wi_sp_12 | 1172064 | 1151017 | 98.2 | 44.62 |
| Wi_sp_13 | 1195862 | 1176887 | 98.4 | 47.19 |
| Wi_sp_14 | 1077166 | 1067151 | 99.1 | 43.82 |
| Wi_sp_15 | 1058848 | 1042823 | 98.5 | 40.49 |
| Wi_sp_16 | 1078208 | 1058635 | 98.2 | 45.33 |
| Wi_sp_17 | 996282 | 984983 | 98.9 | 42.37 |
| Wi_sp_18 | 1097682 | 1088040 | 99.1 | 43.14 |
| Wi_sp_19 | 1121718 | 1113123 | 99.2 | 50.35 |
| Wi_sp_20 | 1077812 | 1058552 | 98.2 | 41.41 |
| Wi_sp_21 | 1143118 | 1132286 | 99.1 | 48.92 |
| Wi_sp_22 | 1087928 | 1061018 | 97.5 | 45.23 |
| Wi_sp_23 | 1120164 | 1111015 | 99.2 | 44.07 |
| Wi_sp_24 | 1192046 | 1173102 | 98.4 | 38.38 |
| ***Rhodnius pallescens*** | **1559728** | **1445663** | **92.7** | **23.27** |
| Rh_pa_01 | 1270496 | 1169917 | 92.1 | 21.99 |
| Rh_pa_02 | 1262894 | 1212213 | 96.0 | 19.64 |
| Rh_pa_03 | 1805108 | 1672486 | 92.7 | 31.37 |
| Rh_pa_04 | 1183370 | 1100466 | 93.0 | 18.20 |
| Rh_pa_05 | 1651000 | 1490111 | 90.3 | 23.92 |
| Rh_pa_06 | 1265832 | 1216668 | 96.1 | 19.59 |
| Rh_pa_07 | 1589572 | 1419243 | 89.3 | 22.54 |
| Rh_pa_08 | 1739134 | 1616983 | 93.0 | 24.67 |
| Rh_pa_09 | 1347828 | 1259470 | 93.4 | 20.20 |
| Rh_pa_10 | 1312996 | 1183628 | 90.1 | 14.49 |
| Rh_pa_11 | 993138 | 875046 | 88.1 | 11.75 |
| Rh_pa_12 | 2515380 | 2354553 | 93.6 | 36.59 |
| Rh_pa_13 | 1910960 | 1807662 | 94.6 | 32.86 |
| Rh_pa_14 | 1835232 | 1727553 | 94.1 | 26.83 |
| Rh_pa_15 | 2102812 | 1938613 | 92.2 | 36.87 |
| Rh_pa_16 | 1169900 | 1085988 | 92.8 | 10.74 |
| **Ixodidae** | **4207274** | **3949263** | **92.2** | **36.31** |
| Am_am_01 | 3133576 | 3030445 | 96.7 | 29.80 |
| Am_am_02 | 7265456 | 6982750 | 96.1 | 55.24 |
| Am_am_03 | 4939078 | 4751748 | 96.2 | 28.81 |
| Am_am_04 | 3386776 | 3298543 | 97.4 | 43.48 |
| Am_am_05 | 4091400 | 3956237 | 96.7 | 44.58 |
| Am_am_06 | 1773478 | 1556865 | 87.8 | 25.62 |
| Am_am_07 | 2235180 | 2154279 | 96.4 | 24.93 |
| Am_ma_01 | 4157520 | 4007437 | 96.4 | 26.50 |
| Am_ma_02 | 5582730 | 5340696 | 95.7 | 49.19 |
| De_va_01 | 5377630 | 5144393 | 95.7 | 30.27 |
| De_va_02 | 4629876 | 4183683 | 90.4 | 36.65 |
| De_va_03 | 5533454 | 5251629 | 94.9 | 46.02 |
| De_va_04 | 5847980 | 5512028 | 94.3 | 49.23 |
| De_va_05 | 1856448 | 1050916 | 56.6 | 19.74 |
| Ix_sc_01 | 2979588 | 2696809 | 90.5 | 21.72 |
| Ix_sc_02 | 4526208 | 4269743 | 94.3 | 49.14 |
