## Supplementary material for "Adapterama III: Quadruple-indexed, double/triple-enzyme RADseq libraries (2RAD/3RAD)": Table S2

**Total number of reads and average coverage after rescuing cut sites with one mismatch and when only the intended cut site (XbaI) is present for *S. tiburo***

Total number of reads obtained from the sequencing runs for each individual of *S. tiburo*, number and percentage of retained reads after cleaning and filtering, and average coverage for the total number of loci in each individual, for both when one mismatch is allowed in the cut site and when only the intended cut site is present. Bold values correspond to averages for the species.

|  | **Raw Reads** | **Ambig. Barcodes** | **Ambig. Radtag** | **Low Quality** | **Retained Reads** | **% Retained Reads** | **Coverage (x)** | **Ambig. Barcodes** | **Ambig. Radtag** | **Low Quality** | **Retained Reads** | **% Retained Reads** | **Coverage (x)** |
| --- | --- | --- | --- | --- | --- | --- | --- | --- | --- | --- | --- | --- | --- |
| ***S. tiburo*** | **1148868** | **63080** | **12630** | **1192** | **1071967** | **93.3** | **17.95** | **63080** | **151962** | **1240** | **796102** | **70.2** | **19** |
| Sp_ti_01 | 1287858 | 50708 | 7033 | 1476 | 1228641 | 95.4 | 18.60 | 50708 | 233786 | 1525 | 778496 | 60.4 | 19.73 |
| Sp_ti_02 | 1308450 | 67622 | 9454 | 1204 | 1230170 | 94.0 | 20.79 | 67622 | 111544 | 1245 | 1031256 | 78.8 | 23.45 |
| Sp_ti_03 | 1191718 | 37732 | 9387 | 1323 | 1143276 | 95.9 | 17.75 | 37732 | 224607 | 1382 | 712114 | 59.8 | 18.71 |
| Sp_ti_04 | 1093198 | 76414 | 6589 | 1077 | 1009118 | 92.3 | 19.98 | 76414 | 67018 | 1095 | 887458 | 81.2 | 21.42 |
| Sp_ti_05 | 1131494 | 44620 | 11182 | 1131 | 1074561 | 95.0 | 17.34 | 44620 | 144525 | 1164 | 813672 | 71.9 | 19.59 |
| Sp_ti_06 | 1107376 | 76964 | 7010 | 1102 | 1022300 | 92.3 | 18.58 | 76964 | 111827 | 1132 | 820750 | 74.1 | 20.37 |
| Sp_ti_07 | 1270442 | 51684 | 12284 | 1617 | 1204857 | 94.8 | 17.90 | 51684 | 252319 | 1675 | 731866 | 57.6 | 18.52 |
| Sp_ti_08 | 1147090 | 70916 | 6366 | 1140 | 1068668 | 93.2 | 20.14 | 70916 | 80472 | 1158 | 922074 | 80.4 | 21.71 |
| Sp_ti_09 | 1386368 | 99576 | 14915 | 1593 | 1270284 | 91.6 | 19.66 | 99576 | 169577 | 1649 | 967076 | 69.8 | 22.37 |
| Sp_ti_10 | 894330 | 41916 | 5880 | 765 | 845769 | 94.6 | 15.91 | 41916 | 100933 | 785 | 663206 | 74.2 | 17.37 |
| Sp_ti_11 | 1309184 | 45526 | 7119 | 1560 | 1254979 | 95.9 | 19.24 | 45526 | 240847 | 1610 | 789474 | 60.3 | 20.19 |
| Sp_ti_12 | 1202502 | 90726 | 6010 | 1156 | 1104610 | 91.9 | 20.85 | 90726 | 84935 | 1179 | 948314 | 78.9 | 22.43 |
| Sp_ti_13 | 1255266 | 50738 | 7458 | 1248 | 1195822 | 95.3 | 20.59 | 50738 | 118741 | 1279 | 979816 | 78.1 | 23.43 |
| Sp_ti_14 | 1434898 | 56612 | 11312 | 1629 | 1365345 | 95.2 | 20.71 | 56612 | 174214 | 1666 | 1046318 | 72.9 | 24.22 |
| Sp_ti_15 | 1248378 | 47730 | 26998 | 1421 | 1172229 | 93.9 | 16.74 | 47730 | 247969 | 1511 | 726690 | 58.2 | 17.79 |
| Sp_ti_16 | 910556 | 60356 | 3976 | 901 | 845323 | 92.8 | 17.16 | 60356 | 50392 | 921 | 752280 | 82.6 | 17.70 |
| Sp_ti_17 | 788708 | 54298 | 31828 | 778 | 701804 | 89.0 | 12.65 | 54298 | 106406 | 839 | 541246 | 68.6 | 13.19 |
| Sp_ti_18 | 953068 | 34390 | 26322 | 1176 | 891180 | 93.5 | 14.28 | 34390 | 196413 | 1254 | 547594 | 57.5 | 15.01 |
| Sp_ti_19 | 1132048 | 69462 | 22399 | 951 | 1039236 | 91.8 | 18.81 | 69462 | 93330 | 1002 | 887620 | 78.4 | 20.35 |
| Sp_ti_20 | 1171024 | 80620 | 23015 | 1155 | 1066234 | 91.1 | 17.73 | 80620 | 135135 | 1211 | 835772 | 71.4 | 19.92 |
| Sp_ti_21 | 785564 | 45996 | 6787 | 733 | 732048 | 93.2 | 14.59 | 45996 | 78395 | 756 | 593000 | 75.5 | 15.58 |
| Sp_ti_22 | 597788 | 38170 | 3698 | 552 | 555368 | 92.9 | 14.39 | 38170 | 28179 | 558 | 505418 | 84.5 | 14.29 |
| Sp_ti_23 | 1479190 | 121934 | 15773 | 1421 | 1340062 | 90.6 | 17.91 | 121934 | 284946 | 1491 | 819366 | 55.4 | 18.57 |
| Sp_ti_24 | 1486342 | 99206 | 20317 | 1495 | 1365324 | 91.9 | 18.55 | 99206 | 310576 | 1665 | 805572 | 54.2 | 18.78 |
