## Supplementary material for "Adapterama III: Quadruple-indexed, double/triple-enzyme RADseq libraries (2RAD/3RAD)": Table S4

**Pairwise F_ST_ values for *S. tiburo* (left) and *S. lewini* (right) populations when allowing a mismatch in the cut site, and when only the intended cut site is present**

Comparison of F_ST_ values in sharks, *S. tiburo* (left) and *S. lewini* (right), obtained when both, we allowed mismatches in the cut site permitting the third enzyme be present in the 3RAD data, and when only the exact intended cut sites are present in the 3RAD. Values are similar varying only in the last decimal position in both datasets.

|  | **TB** | **CK** | **NAFL** | **CAFL** | **TAB** | Rescuing Cut Sites | *Sphyrna tiburo* |
| --- | --- | --- | --- | --- | --- | --- | --- |
| **CAM** | 0.082 | 0.089 | 0.235 | 0.234 | 0.071 |  |  |
| **TB-CH** | | 0.046 | 0.073 | 0.073 | 0.051 |  |  |
| **CK** |  |  | 0.083 | 0.083 | 0.051 |  |  |
| **NAFL** |  |  |  | 0.364 | 0.080 |  |  |
| **CAFL** |  |  |  |  | 0.080 |  |  |
|  | **TB** | **CK** | **NAFL** | **CAFL** | **TAB** | Intended Cutsite |  |
| **CAM** | 0.081 | 0.086 | 0.235 | 0.232 | 0.071 |  |  |
| **TB-CH** | | 0.046 | 0.073 | 0.072 | 0.049 |  |  |
| **CK** |  |  | 0.083 | 0.082 | 0.051 |  |  |
| **NAFL** |  |  |  | 0.366 | 0.079 |  |  |
| **CAFL** |  |  |  |  | 0.078 |  |  |

|  | **C** | **LC** | **PM** | **SC** | **T** | **VC** |  | *Sphyrna lewini* |
| --- | --- | --- | --- | --- | --- | --- | --- | --- |
| **BoC** | 0.155 | 0.229 | 0.218 | 0.122 | 0.119 | 0.125 | Rescuing Cut Sites |  |
| **C** |  | 0.222 | 0.212 | 0.120 | 0.118 | 0.123 |  |  |
| **LC** |  |  | 0.381 | 0.155 | 0.161 | 0.167 |  |  |
| **PM** |  |  |  | 0.155 | 0.157 | 0.165 |  |  |
| **SC** |  |  |  |  | 0.097 | 0.102 |  |  |
| **T** |  |  |  |  |  | 0.097 |  |  |
|  | **C** | **LC** | **PM** | **SC** | **T** | **VC** |  |  |
| **BoC** | 0.155 | 0.228 | 0.219 | 0.122 | 0.120 | 0.124 | Intended Cut Site |  |
| **C** |  | 0.222 | 0.214 | 0.121 | 0.117 | 0.121 |  |  |
| **LC** |  |  | 0.378 | 0.156 | 0.161 | 0.167 |  |  |
| **PM** |  |  |  | 0.157 | 0.158 | 0.165 |  |  |
| **SC** |  |  |  |  | 0.097 | 0.101 |  |  |
| **T** |  |  |  |  |  | 0.097 |  |  |
