## Supplementary figures and images for "Adapterama III: Quadruple-indexed, double/triple-enzyme RADseq libraries (2RAD/3RAD)"

### Fig. S1

#### 3RAD Quadruple-Indexed Libraries

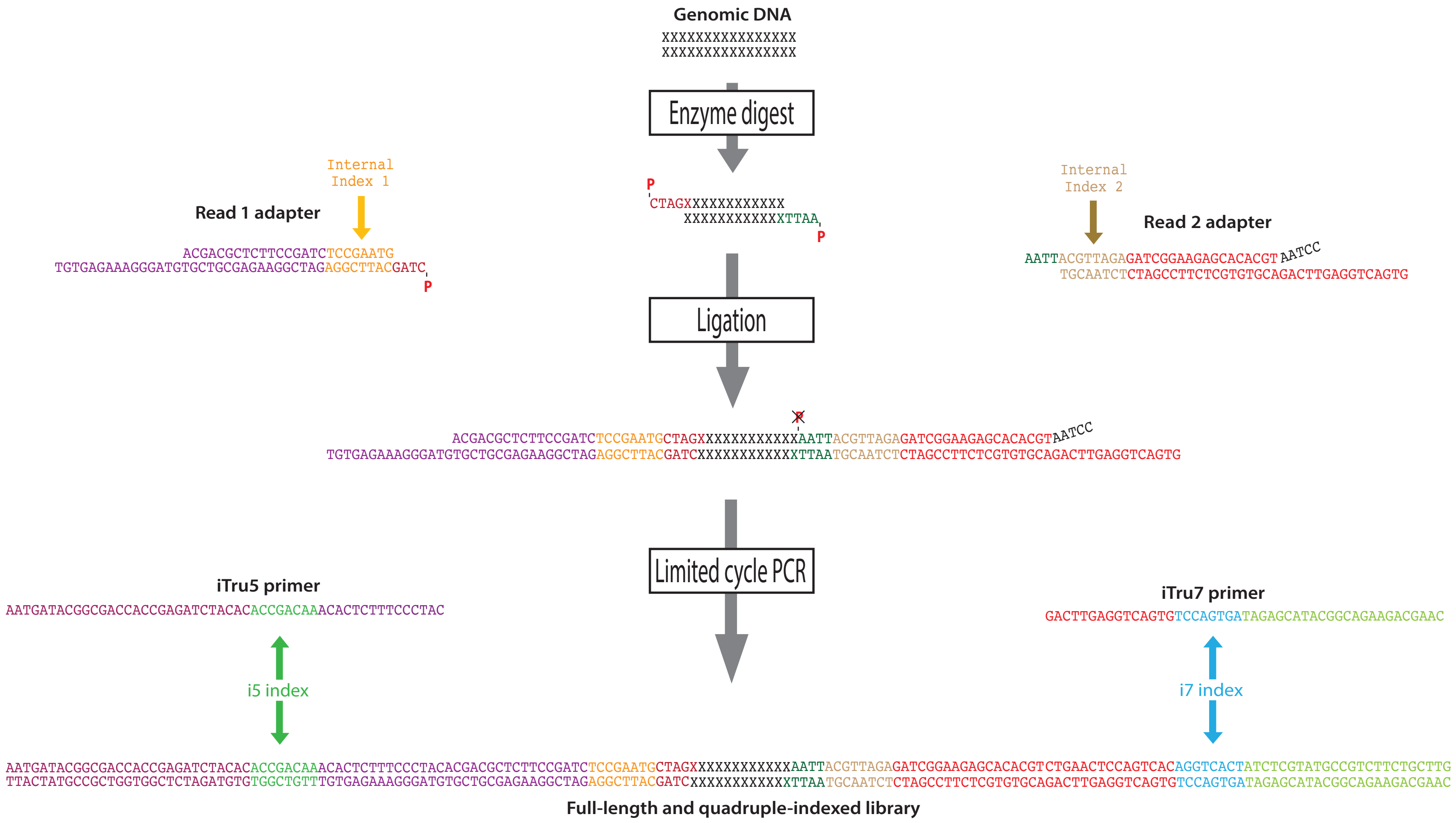

### Fig. S2

#### Quadruple-Indexed 3RAD Libraries

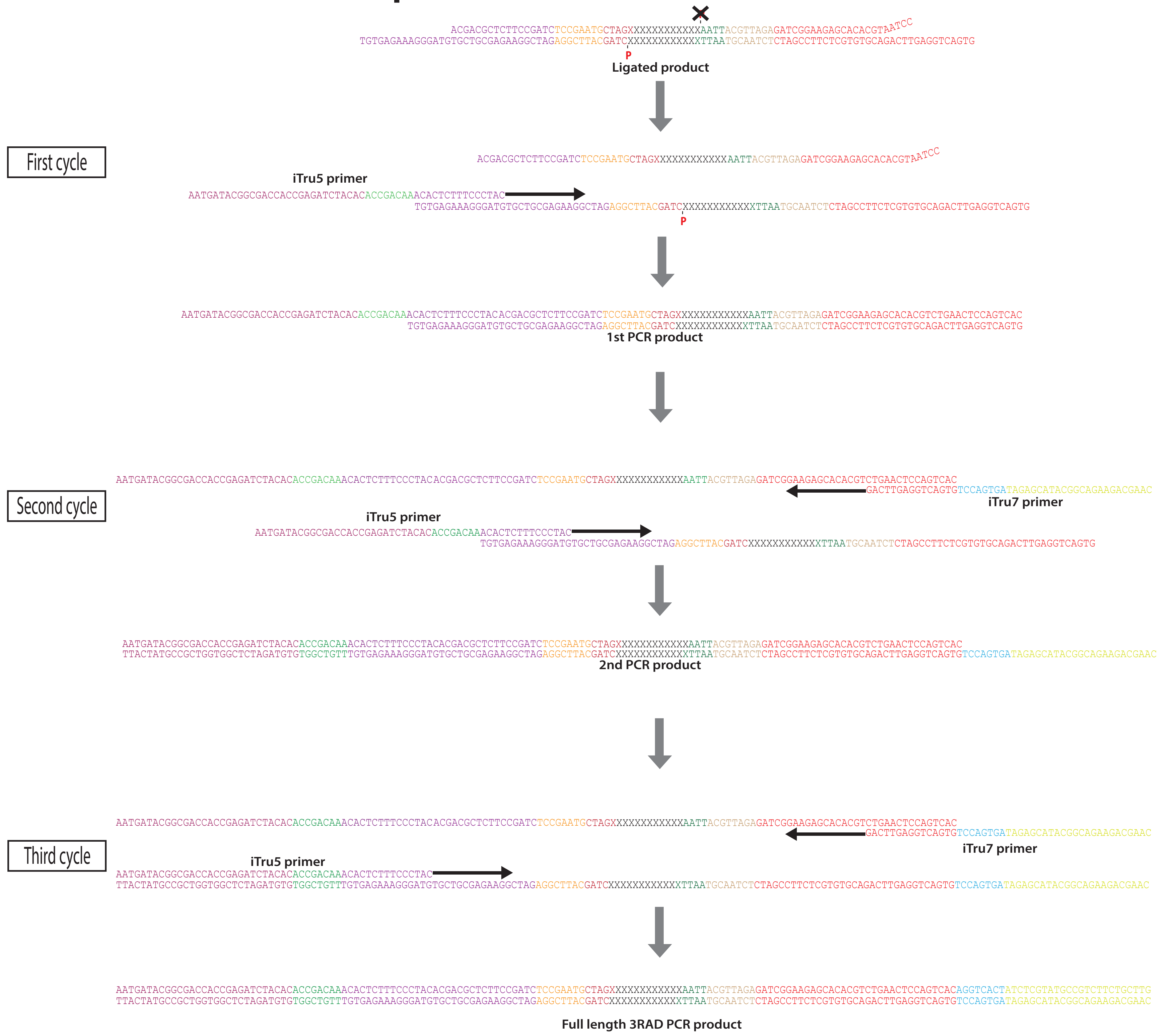

### Fig. S3

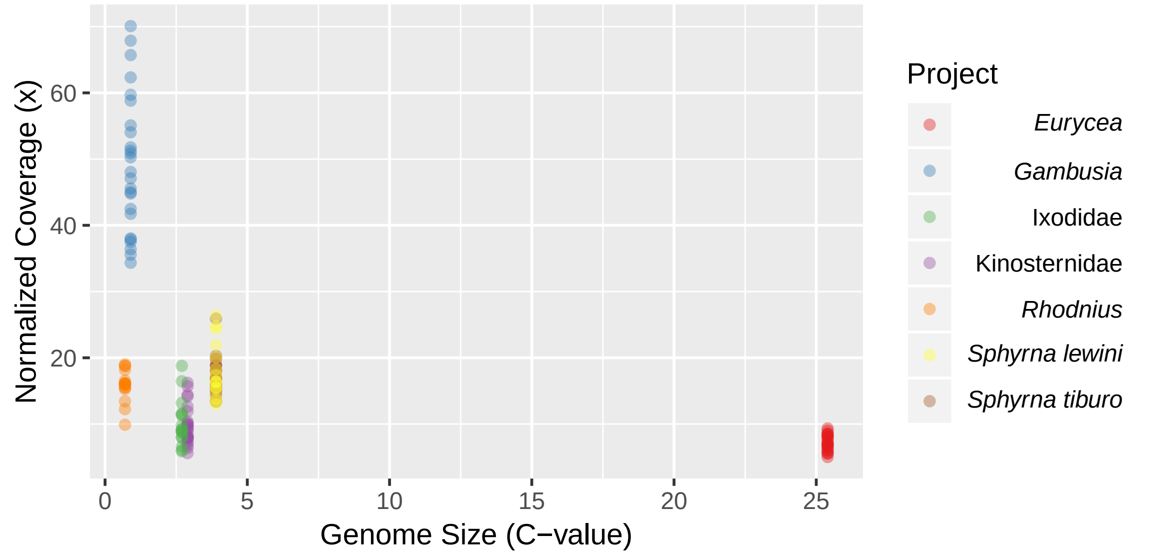

### Fig. S4

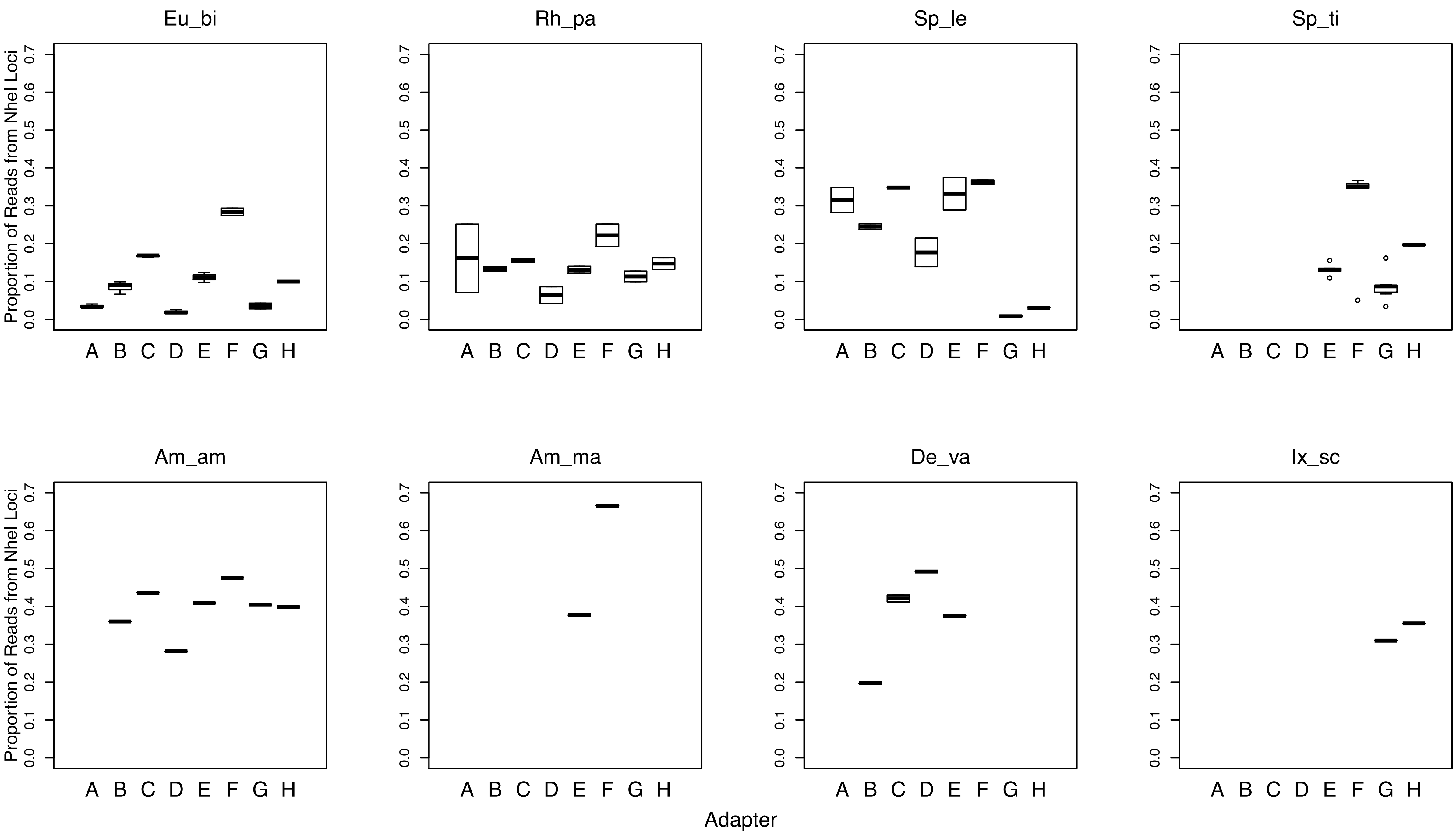
