## Supplementary material for "Adapterama III: Quadruple-indexed, double/triple-enzyme RADseq libraries (2RAD/3RAD)": File S1

**High-Throughput 3RAD Protocol**

1. Normalize all DNA samples to 20 ng/µL (quantified on Qubit), placing ≥ 20 µL in plates or strip tubes. Ensure these (and all other tubes/plates below) are labeled clearly. Protocol will work with much less DNA, but ≥ 5 ng/µL is best.
2. Set up digestion with the following per-sample recipe:
   1.5 μL NEB 10x CutSmart Buffer
   5.0 μL dH_2_O
   0.5 µL XbaI [20.000 U/mL] (or another Read 1 enzyme)
   0.5 µL EcoRI-HF [20.000 U/mL](or another Read 2 enzyme)
   0.5 μL NheI [20.000 U/mL] (or another Read 1 adapter dimer-cutting enzyme)
   1.0 µL ds iTru NheI adapter [5 µM] (or another suitable adapter; see Table 2)
   1.0 µL ds iTru EcoRI adapter [5 µM] (or another suitable adapter; see Table 2)
   5.0 µL genomic DNA (20 ng/µL; if dilute, use a greater volume & reduce water)
3. Incubate samples at 37°C for 1 hr. in a thermal cycler. Proceed immediately to Step 4.
4. Add the ligation reagents. We recommend preparing a master mix shortly before the digestion is completed and adding the following per-reaction mix (i.e., 5 μL) to each tube:

2.0 μL dH_2_O
1.5 µL ATP (10 mM); note: rATP, NOT dATP
0.5 µL 10x Ligase Buffer (ensure components are in solution [warm it up!])
1.0 µL DNA T4 ligase [100 units/µL; dilute from 400 units/µL stock using	Diluent Buffer A]

5. Incubate at 22°C for 20 min., 37°C for 10 min., 22°C for 20 min., 37°C for 10 min., 80°C for 20 min., then hold at 10°C.
6. If the multiple ligation products that you have at this step share adapter combinations and therefore pooling is not an option, please continue to step 7).

Pooling ligation products that have unique adapter combinations (from Step 2)

a. Using the multi-channel pipette and changing tips each time, skloosh, then take 10 µL from each ligation and add it to a strip tube. Thus, when using a 96-well plate, each tube in the strip will have 120 µL. Seal the plate, label it well, and freeze it for potential use later.
b. Pooling step 2: Into a single 1.5 mL tube, pool 60 µL from each tube of the strip from Step 6a. This should yield 480 µL of ligation product into the 1.5 mL tube. Label the strip tube well and freeze it for potential future use.
c. Split the pooled ligation products into two 1.5 mL tubes (measured as 2 x 120 µL).

7. Clean and purify ligation products. Either:

For individual ligation products

To the 20 µL of ligation product add 30 µL of dH_2_O, then add 60 µL of SpeedBeads (Thermo-Scientific, Waltham, MA, USA), purify as normal and resuspend in 20 µL of dH_2_O.

**OR**

From pooled ligation products:
To each ligation pool of 240 µL then add 300 µL of SpeedBeads (i.e. 1.25x speedbeads; measured as 2 x 150 µL). Purify as normal and resuspend each in 30 µL of dH_2_O, then combine into one tube (total of 60 µL).

Set up PCR reactions. Either:

PCR recipe to add complete Illumina adapters & indexes in **individual** cleaned-ligation products:

5.0 μL 5x Kapa HiFi Buffer
0.75 μL dNTP’s (10 mM stock from Kapa kit)
8.75 μL dH_2_O (to make final total volume 50µL)
0.5 μL Kapa HiFi DNA Polymerase (1 unit/μL from Kapa kit)
2.5 µL i5 Primer (5µM -> 0.5 µM final)
2.5 µL i7 Primer (5µM -> 0.5 µM final)
5 μL Linker ligated-cleaned DNA fragments from Step 7 (placed on magnet).

**OR**

PCR recipe to add complete Illumina adapters & indexes in **pooled** ligation products (three replicates per pool):

10.0 μL 5x Kapa HiFi Buffer (Kapa Biosystems, Wilmington, MA)
1.5 μL dNTP’s (10 mM stock from Kapa kit)
7.5 μL dH_2_O (to make final total volume 50µL)
1.0 μL Kapa HiFi DNA Polymerase (1 unit/μL from Kapa kit)
5.0 µL i5 Primer (5µM -> 0.5 µM final)
5.0 µL i7 Primer (5µM -> 0.5 µM final)
20.0 μL Linker ligated DNA fragments from step 7 (placed on magnet).

Cycling for both methods: 98°C for 1 min.; then, 12 cycles of: 98°C for 20 sec., 60°C for 15 sec., 72°C for 30 sec.; 72°C for 5 min. Hold at 15°C.

8. Purify with SpeedBeads by either:

To **individual** PCR reactions
Add 30 µL of dH_2_O, + 60 µL SpeedBeads, skloosh; purify as normal, and resuspend in 20 µL dH_2_O, incubate few minutes at room temperature, then place on magnet and pull off all liquid (~18 µL), leaving the beads behind.

**OR**

To each of the **pooled-ligations** PCR reactions
Add 50 µL of dH_2_O, + 100 µL SpeedBeads, skloosh; purify as normal, and resuspend in 20 µL dH_2_O. Pool beads from all 3 PCR replicates (60 µL total), then place on magnet and pull off all liquid (~55 µL), leaving the beads behind.

9. Run 5 µL on agarose gel to ensure each sample/pool worked.
10. Quantify with Qubit, normalize, pool (those that were not pooled after ligation), SpeedBead (1:1.25, dna:speedbead), and size select on Pippin (525 bp +/- 10%). [change size as necessary to avoid bright bands; we do *not* know that choosing a narrow size-range is best]
11. Quantify with Qubit, mix in appropriate proportions with other libraries, and send off to NovaSeq, HiSeq, NextSeq or MiSeq: Use TruSeq primers & do dual (8nt) index reads!
