## Supplementary material for "Adapterama III: Quadruple-indexed, double/triple-enzyme RADseq libraries (2RAD/3RAD)": File S4

How to Handle Plates with 3RAD v2 Adapter Aliquots

When you receive the adapters, there is 1.625 nmol of each oligo pair dried in each well, but they are **NOT** annealed. You will need to reconstitute them to the appropriate volume (64 µL -> 25 µM), anneal them, then dilute again (to 5 µM) & aliquot them.

Liquid for reconstitution & annealing (10 mM Tris pH 8, 0.1 mM EDTA, 100 mM NaCl):

For 50 mL of salty TLE, add the following to a 50mL conical:

40 mL dH_2_O

500 µL 1M Tris pH 7.5 to 8

20 µL 0.5M EDTA pH 8

1 mL of 5 M NaCl (or 5 mL of 1M NaCl)

Fill with distilled water to 50 mL mark.

**Protocol:**

1) Centrifuge the dry plates to get all the adapter to the bottom of the wells.

2) To limit contamination, peel back the foil cover from the plate one row at a time to reconstitute.

3) Add 64 µL of the liquid from above to each well.

-Use the pipet tip to help scrap the bottom of the well to dislodge any of the adapters that is stuck.

-Skloosh (pipette up & down) several times to mix.

-Wait a few minutes.

- Skloosh several more times to mix.

-Let the adapters sit in the liquid at room temperature for at least 5 minutes.

The adapters are now at 25 µM.

4) Anneal the adapters together:

-Use thermocycler to denature (95°C for 1 min.) & cool slowly (e.g., 0.1°C per sec.).

5) Dilute aliquots of the annealed adapters into new labeled strip tubes:

- Add 10 µL of annealed adapters to 40 µL of salty TLE (final conc. = 5 µM), for SIX separate strips. [Note: This will leave ~4 µL behind in the oligo plates (could be worth getting as much as possible for the Read1 Adapters – they are limiting; but the Read2’s are in excess, so leave that behind). Each strip (with 50 µL) can do FOUR full plates.]

6) Store the strip tubes of adapters at -20°C.

7) Before each use, take adapter aliquots out to thaw and **skloosh well before using!**
