## Supplementary material for "Adapterama III: Quadruple-indexed, double/triple-enzyme RADseq libraries (2RAD/3RAD)": File S5

**Supplementary methods and results for 3RAD efficiency with low-quantity samples: a proof of concept**

We tested and compared our 3RAD protocol with the traditional ddRAD protocol (using the two flanking REs from 3RAD and heat-killing after ligation) and 2RAD (using the two flanking REs from 3RAD, but no heat-killing after ligation) with an array of sample dilutions.

To simplify the comparison between protocols, we used as template the pUC19 vector, which contains XbaI cut-site at position 423 and EcoRI cut-site at position 396. Our methods were as follows:

1. Amplified a ~500 bp fragment within the vector using the primers pUC19-215F-AAGGAGAAAATACCGCATCAGG and PUC19-774R-TAACCGTATTACCGCCTTTGAG and the following PCR recipe and conditions.

5.0 μL 5x Kapa HiFi Buffer
1.0 μL dNTP’s (10 mM stock from Kapa kit)
33 μL dH_2_O (to make final total volume 50µL)
2.0 μL Kapa HiFi DNA Polymerase (1 unit/μL from Kapa kit)
5.0 µL Forward Primer (5µM -> 0.5 µM final)
5.0 µL Reverse Primer (5µM -> 0.5 µM final)

3.3 μL input DNA

Cycling conditions: 95°C for 3 minutes (min), then 28 cycles of 95°C for 30 sec, 50°C for 30 sec, 72°C for 30 sec; and a final extension at 72°C for 5 minutes followed by holding at 4°C.

2. Purified the PCR product with SpeedBeads with a 1:1 ratio and quantified with a Qubit fluorometer. This measured 15 ng/μL.
3. Made a 10-fold dilution series. We used these six products (one stock and five dilutions) as input for the 3RAD, 2RAD and ddRAD libraries.
4. Digested with REs using the following recipe:

|  | **3RAD** | **2RAD** | **ddRAD** | **Negative** |
| --- | --- | --- | --- | --- |
| **CutSmart** | 1.5μL | 1.5 μL | 1.5 μL | 1.5 μL |
| **Adapters (5 μM**) | 1 μL/ each | 1 μL/ each | 1 μL/ each | 1 μL/ each |
| **DNA** | 3.3 μL | 3.3 μL | 3.3 μL | 3.3 μL |
| **Water** | 6.7 μL | 7.2 μL | 7.2 μL | 10.5 μL |
| **XbaI (20,000 U/mL)** | 0.5 μL | 0.5 μL | 0.5 μL | 0.5 μL |
| **EcoRI-HF (20,000 U/mL)** | 0.5 μL | 0.5 μL | 0.5 μL | 0.5 μL |
| **NheI (20,000 U/mL)** | 0.5 μL | 0 | 0 | 0 |

Cycling conditions: 37°C for 1 hour. For the ddRAD experiment, we heat-killed REs with a 20 min incubation at 65°C following digestion.

5. Immediately added the ligation mix (10 μL) to each sample. That mix was as follows:

   7.25 μL dH_2_O
   1.5 µL ATP (10 mM); note: rATP, NOT dATP
   1.0 µL 10x Ligase Buffer (ensure components are in solution [warm it up!])
   0.25 µL DNA ligase 400 units/µL

   Cycling conditions: 22 °C for 20 min., 37 °C for 10 min., 22 °C for 20 min., 37 °C for 10 min., 80 °C for 20 min., then hold at 10 °C.
6. Set up PCR reactions with the following recipe per-sample:

5.0 μL 5x Kapa HiFi Buffer
0.75 μL dNTP’s (10 mM stock from Kapa kit)
3.75 μL dH_2_O (to make final total volume 50µL)
0.5 μL Kapa HiFi DNA Polymerase (1 unit/μL from Kapa kit)
2.5 µL i5 Primer (5µM -> 0.5 µM final)
2.5 µL i7 Primer (5µM -> 0.5 µM final)
10.0 μL input DNA from Step 5

Cycling conditions: 95 °C for 2 min, followed by 15 cycles of 98 °C for 20 sec, 60 °C for 30 sec and 72 °C for 30 sec; and a final extension at 72 °C for 5 min, holding after at 12 °C.

7. Ran the PCR products of each of the three experiments on a 1.5% agarose gel. For every DNA input concentration tested, the first well corresponds to 3RAD, the second to 2RAD, and the third to ddRAD library.

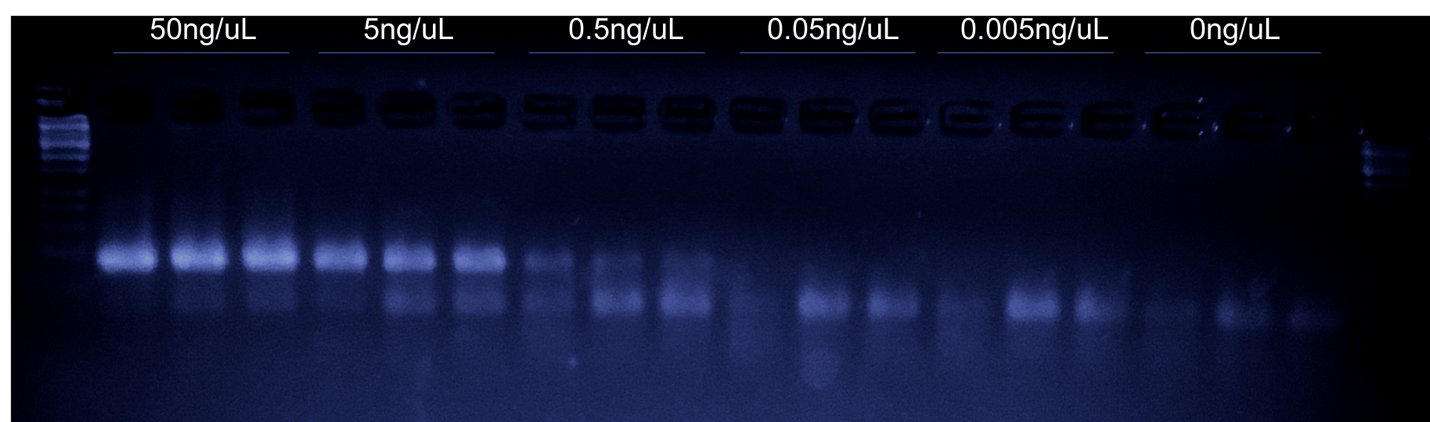
