## Supplementary material for "Adapterama III: Quadruple-indexed, double/triple-enzyme RADseq libraries (2RAD/3RAD)": File S6

**Supplementary methods and results for 3RAD example datasets**

***Guidelines for library preparation in all projects***

We digested sample DNA for 1 hr at 37 °C in a reaction mix that consisted of: 1.5 µL 10x CutSmart Buffer, 5.0 µL dH_2_O, 0.5 µL of XbaI at 20 U/µL, 0.5 µL of EcoRI-HF at 20 U/µL, 0.5 µL of NheI at 20 U/µL, 1 µL 5 µM double-stranded iTru R1.A adapter, 1 µL 5 µM double-stranded iTru R2.1 adapter, and 5 µL DNA. After incubation at 37 °C for 1 hour, we added 2.0 µL dH_2_O, 1.5 µL ATP (10 µM), 0.5 µL 10x Ligase Buffer, and 1.0 µL T4 DNA Ligase (100 units/µL, NEB M0202L buffer diluted 1:3 in NEB B8001S enzyme dilution buffer) to each reaction, and we digested/adapter-ligated mixtures in a thermocycler with the following conditions: 22 °C for 20 min. and 37 °C for 10 min. for two cycles followed by a single cycle of 80°C for 20 min. After ligation, we pooled all *Wisteria*, *Gambusia*, and Kinosternidae individuals by project; we maintained *Eurycea,* *Rhodnius*, *Sphyrna*, and Ixodidae samples individually. We proceeded immediately to a pre-PCR clean-up (File S1) to remove remaining reagents and unincorporated adapters, using Sera-Mag SpeedBeads (Thermo Fisher Scientific, Waltham, MA, USA; see Glenn et al., 2016 for preparation methods) at a ratio of 1.2:1 SpeedBeads to DNA, resuspending DNA in 20 µL of TLE.

To generate full-length library constructs, we conducted a PCR with 5.0 µL of this cleaned, post-ligation DNA, 5.0 µL Kapa HiFi Buffer (Roche, Basel, Switzerland), 0.75 µL dNTPs (10 mM), 8.75 µL dH_2_O, 0.5 µL Kapa HiFi DNA Polymerase (1 unit/µL), 2.5 µL iTru5 primer (5 µM), and 2.5 µL iTru7 primer (5 µM) and thermocycler conditions as follows: 95°C for 2 min.; then, 14 cycles of 98°C for 20 sec., 60°C for 15 sec., 72°C for 30 sec.; 72°C for 5 min.; hold at 15°C. To validate that the library preparation process was successful, we ran 5 µL of PCR product with 2 µL loading dye on a 1.5% agarose gel for 45 minutes at 90 volts along with Hi-Lo DNA Marker (Minnesota Molecular, Minneapolis, MN, USA). A smear of evenly distributed and bright DNA around ~300-800 bp, without noticeable bands in this target size zone, indicated successful library preparation. After validation, we cleaned the remaining PCR reaction volume with SpeedBeads in at least 1:1 (SpeedBeads:DNA) ratio, and we eluted cleaned DNA in 20 µL of TLE.

We quantified cleaned libraries using either a Qubit Fluorimeter (Thermo Fisher Scientific, Waltham, MA, USA) or SYBR Green assay with a plate reader, and for those projects with samples not already pooled, we pooled equal amounts prior to size-selection. Pools contained up to 96 samples and/or totaled 1-1.8 µg. We size-selected pooled libraries using a Pippin Prep (Sage Science, Beverly, MA, USA) with a 1.5% dye-free Marker K agarose gel cassette (CDF1510) set to capture fragments at 550 bp +/-10%. If < 1 ng/µL of DNA was recovered following size-selection, we further amplified libraries with P5 and P7 primers (Glenn et al., 2016) with 6-12 cycles of 98°C for 20 sec., 60°C for 15 sec., 72°C for 30 sec.; followed by 72°C for 5 min. and holding at 15°C. We cleaned the PCR reaction with SpeedBeads using a ratio ≥ 1:1 (SpeedBeads:DNA), and we eluted cleaned DNA in 25 µL of TLE.

Data from each project were assembled independently using Stacks v1.40, v1.42, or v.1.44 (Catchen et al*.*, 2013; Catchen et al*.*, 2011). We used the *process_radtags* program to demultiplex and/or clean and trim the sequence data. We removed reads with an uncalled bases (-c) and discarded reads with low quality scores (-q) with a default sliding window of 15% of the length of the read and raw Phred score of 10. We specified XbaI and EcoRI as REs, and we rescued sequence tags (internal tags) and RAD-tags (enzyme over-hang) within 2 mismatches of their expected sequence (-r); otherwise, reads were discarded. We truncated (-t) PE150 reads to 140 nt and PE75 reads to 64 nt to have equal length among all reads with different barcodes.

We parallel-merged the mates of paired-end reads using the ‘paste -d’ Unix command. We used the Stacks *denovo_map* pipeline for each project with the following settings: the minimum number of identical raw reads required to create a stack (-m) = 3, the maximum distance between stacks (-M) = 3, and the number of mismatches allowed between samples when generating the catalog (-n) = 4. Coverage, number of loci, and number of SNPs recovered were scored for each species and compared to genome size and sequencing read length (PE75 or PE150). Next, we used the Stacks *populations* program with the following settings: the minimum populations per locus (-p) = 60–75% of total populations and the minimum individuals (within a population) per locus (-r) = 75%.

For projects with predefined populations or collection localities, we calculated F-statistics in Stacks. For all datasets, we performed preliminary Structure v2.3.4 (Pritchard, Stephens, & Donelly, 2000) analyses using burn-in and sampling lengths of 10,000 and 100,000 MCMC repeats, respectively. We tested values of *K* from 1 to the number of predefined populations, or in the case of projects without predefined populations, we tested values of *K* as described below. We did this using an admixture model with correlated allele frequencies and with three iterations for each value of *K*. We chose a value of *K* with the highest mean likelihood across iterations and plotted the iteration with the highest likelihood for that *K*. We present these results, not as definitive answers to biological questions involving these taxa, but as examples to illustrate 3RAD data. For detailed information about assembly results (e.g., raw reads, coverage, etc.), see File S1.

***Project: Kinosternidae***

In order to show the utility of 3RAD across multiple evolutionary time scales, we used a sampling approach that would allow us to simultaneously gather data for both higher-level systematics-based and population genetics analyses. Thus, we sampled four individuals each from the three major river drainages within the distribution of *Sternotherus depressus* (12 total turtles) and two individuals each from the remaining species of *Sternotherus* (*S. peltifer, S. minor, S. carinatus, S. odoratus*), along with two individuals each of the more distantly related *Kinosternon baurii* and *Staurotypus triporcatus* which allowed our data set to span the root node of Kinosternidae.

We used *process_radtags* as described above to demultiplex the individuals. Due to the relatively old crown age (ca. 55my; Iverson, Le, & Ingram, 2013) and phylogenetic context we were using, we chose to assemble data *de novo* with pyRAD-3.0.4 (Eaton, 2014) on the University of Alabama’s RC2 high performance computing cluster. We set the within- and between- cluster threshold to 88% similarity and minimum within-individual sequence depth to ≥8. We generated three data matrices: 1) 10% missing data within crown *Sternotherus* (ca 15my; Iverson, Le, & Ingram, 2013) and a greater amount of missing data for more distantly related individuals; 2) 20% missing data across the entire dataset; and, 3) 34% missing data across the entire dataset. These final matrices included: 1) 6,051; 2) 4,034 (Table 3 in main text); and, 3) 8,312 loci respectively. We inferred phylogenetic relationships among all sampled turtles from the concatenated dataset using maximum likelihood (ML) inference in RAxML 8.2.8 (Stamatakis, 2014) and the GTRGAMMA model for sequence evolution. We estimated branch support for each node using the automatic bootstrap function on the CIPRES science gateway (Pattengale et al., 2010; Miller, Pfeiffer, & Schwartz, 2010), which uses a stopping rule to calculate when sufficient replications have been completed.

Phylogenies inferred from all three datasets recovered the same relationships with high support (bootstrap = 99–100) for all nodes, except the root of *Sternotherus* (Fig. 1). Poor support at this node is not unsurprising, as ongoing work (Scott, Glenn, & Rissler, 2017) show support for rapid diversification within crown *Sternotherus*.

To assess the utility of these data for population-level inference, we separately analyzed the twelve *Sternotherus depressus* samples collected from three river drainages (i.e., four individuals each). We assembled these data using the Stacks pipeline as described above. We stipulated a 75% (-p) complete matrix, and we recovered 16,695 loci. Pairwise F_ST_ values ranged from 0.109 and 0.148 (Table 1), matching expectations and Structure results (Fig. 2).

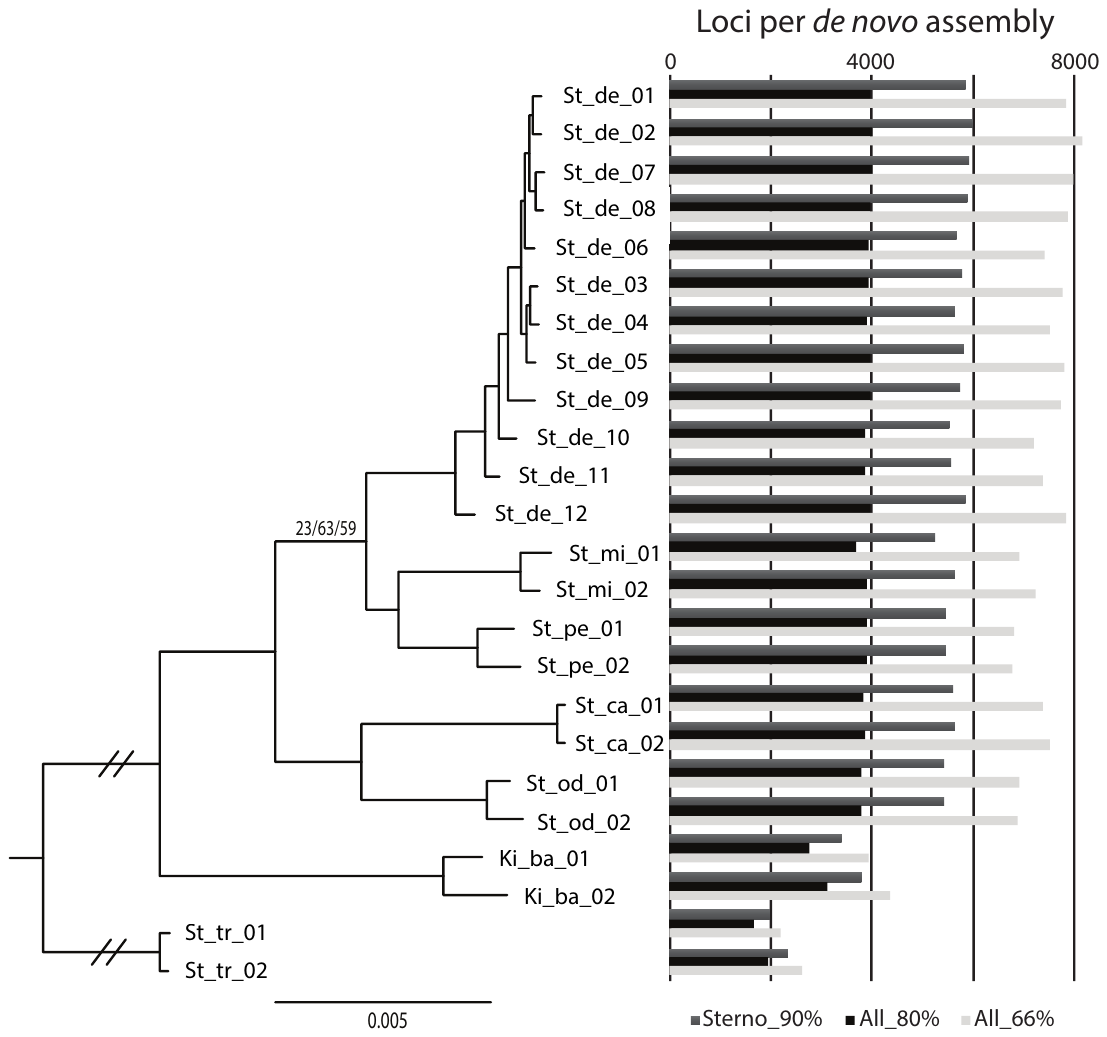

**Figure 1. Maximum likelihood phylogeny of Kinosternidae from 3RAD loci**

All major nodes are supported with bootstrap values ≥ 99 except where noted. On the right, a bar plot shows the number of loci recovered in *de novo* assemblies with different amounts of missing data. Sterno_90% = loci present in ≥ 90% *Sternotherus* individuals with greater missing data allowed for other taxa; All_80% = loci present in ≥ 80% of individuals; All_66% = loci present in ≥ 66% of individuals.

**Table 1. Pairwise F_ST_ values for *Sternotherus depressus* from three localities**

Estimated from an assembly of 25,578 SNPs from 16,695 3RAD loci.

|  | **Mulberry** | **North** |
| --- | --- | --- |
| **Sipsey** | 0.109 | 0.148 |
| **Mulberry** |  | 0.144 |

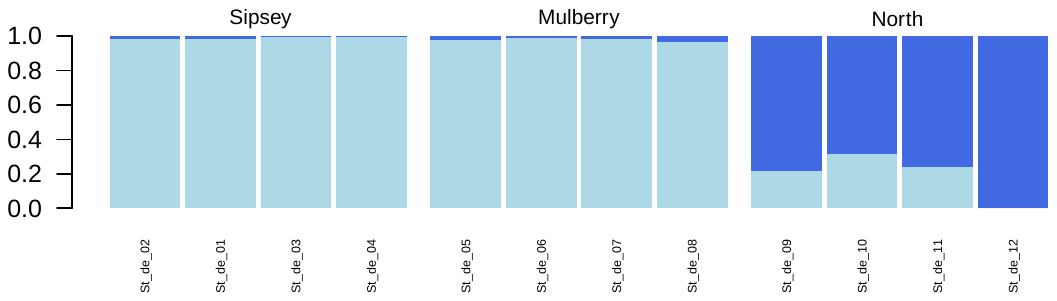

**Figure 2.** **Structure plot for *Sternotherus depressus* (Kinosternidae) with K = 2**

Estimated from an assembly of 25,578 SNPs from 16,695 3RAD loci. Predefined populations are separated by large spaces and labeled above bars; individual sample names are shown below bars.

***Project: Ixodidae***

We included samples from four tick species of the Family Ixodidae, which includes vectors of several diseases such as human monocytotropic ehrlichiosis, tularemia, and Lyme disease. We constructed 3RAD libraries from seven *Amblyomma americanum* samples, two *A. maculatum* samples, five *Dermacentor variabilis* samples and two *Ixodes scapularis* samples following the independent PCR (i.e., repetitive inner tags) protocol (Figure 3 in the main text; File S1). We used the lone star tick (*A. americanum*) dataset, with samples collected from two localities in the United States, for population-level analyses. We collected samples in August of 2015 from hunters at deer check stations in Stanley County, North Carolina and Johnson and Adair Counties, Kentucky. We tested values of *K* = 1 and *K* = 2 in Structure.

We assembled 69,518 SNPs from 19,843 3RAD loci and calculated a pairwise F_ST_ value of 0.407 between samples from North Carolina and Kentucky. Despite this high F_ST_, our Structure analyses supported *K* = 1 (Fig. 3). Notably, other authors have also suggested a lack of population structure in this species at a local scale and have attributed this to dispersal on animal hosts and rapid population expansions (Mixson et al., 2006; Trout, Steelman, & Szalanski, 2010). Greater attention should be given to understanding the population structure of this species, a vector of multiple human pathogens.

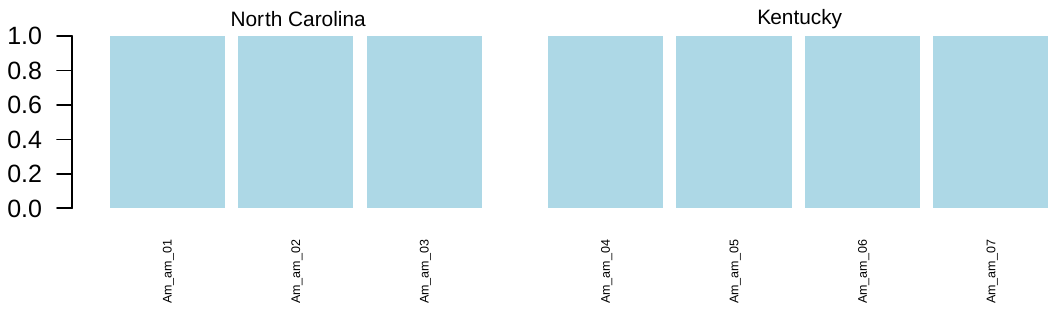

**Figure 3. Structure plot for *Amblyomma americanum* (Ixodidae) with K = 1**

Estimated from an assembly of 69,518 SNPs from 19,843 3RAD loci. Predefined populations are separated by large spaces and labeled above bars; individual sample names are shown below bars.

***Project:* Eurycea bislineata *species complex***

We included 21 individuals representing geographically widespread and representative samples from all five named taxa of the two-lined salamander (*Eurycea bislineata*) species complex. We prepared these libraries and analyzed these data using the workflow described above. We tested values from *K* = 1 to *K* = 10.

Predictably, due to the large size of the genome of these salamanders, we obtained relatively low-coverage data. While we are using these low-coverage data for downstream phylogenetic analyses, here, we report only basic statistics regarding the number and coverage of loci and SNPs recovered. Our Structure analysis supported *K* = 5, with these groups in concordance with some evolutionary relationships suggested by previous allozyme and mitochondrial data (Jacobs, 1986; Kozak, Blaine, & Larson, 2006; Fig. 4).

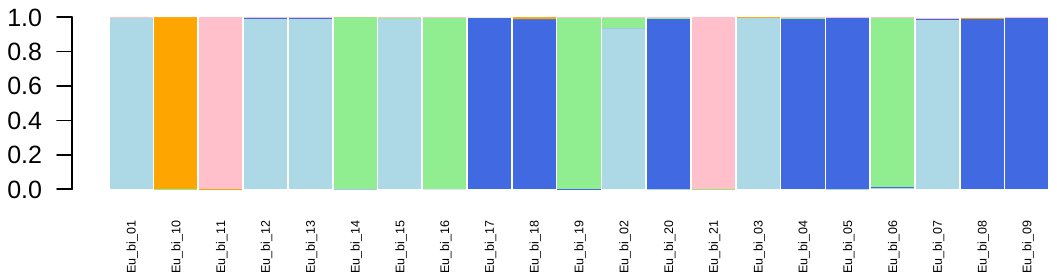

**Figure 4. Structure plot for the *Eurycea bislineata* species complex with K = 5**

Estimated from an assembly of 360 SNPs from 30 3RAD loci. Individual sample names are shown below bars.

***Project:* Wisteria**

We included twenty-four samples of the invasive vine, *Wisteria* spp., collected in Athens, Georgia, USA. We used a slightly modified 3RAD protocol (Supplemental File S7). We used 0.166 µL of the restriction enzyme NheI-HF per reaction in the digestion, instead of 0.5 µL. After ligation, we pooled 7 µL from all samples and immediately cleaned this pool with SpeedBeads using a 1:1 SpeedBeads:DNA ratio, using two washes of 80% EtOH and eluting into 33 µL TLE. In six 50 µL PCR reactions, we combined 5 µL of pooled DNA, 5 µL of 5 µM iTru5-8N (Hoffberg *et al.* 2016), 1.5 µL dNTPs at 10 mM, 10 µL KAPA HiFi Buffer, and 1 µL of KAPA HiFi Hotstart DNA Polymerase. Thermocycler conditions were as follows: 95°C for 2 minutes, 98°C for 20 sec, 61°C for 15 sec, 72°C for 30 sec, and 72°C for 5 min. We pooled these PCR products and cleaned them using a 2:1 SpeedBeads:DNA ratio. In four 50 µL PCR reactions, we combined 5 µL of pooled DNA, 5 µL of 5 µM P5, 5 µL of 5 µM iTru7, 1.5 µL of dNTP at 10 mM, 10 µL KAPA HiFi Fidelity Buffer, and 1 µL of KAPA HiFi Hotstart DNA Polymerase. Thermocycler conditions were as follows: 95°C for 2 min; 6 cycles of 98°C for 20 sec, 61°C for 15 sec, 72°C for 30 sec; and 72°C for 5 min. We cleaned the four PCR reactions pool with SpeedBeads, quantified it with a Qubit, and size-selected as described above. We conducted a final PCR with P5/P7 primers to increase library concentration. In three 50 µL reactions, we combined 10 µL of DNA, 5 µL of 5 µM of each primer, 1.5 µL of dNTPs at 10 mM, 10 µL KAPA HiFi Fidelity Buffer, and 1 µL of KAPA HiFi Hotstart DNA Polymerase. Thermocycler conditions were as follows: 95°C for 2 min; 18 cycles of 98°C for 20 sec, 61°C for 15 sec, 72°C for 45 sec; and 72°C for 5 min. We pooled PCR products, cleaned them with a 2:1 SpeedBeads:DNA ratio, quantified them, and combined them with unrelated libraries for sequencing on an Illumina NextSeq High Output v2 150 cycle kit to yield PE75 reads. We assembled data as described above. We tested values from *K* = 1 to *K* = 3.

Because all *Wisteria* samples originated from the same population in Athens, GA, we did not calculate F_ST_ values. Our Structure analysis supported *K* = 2, suggesting ancestry from two groups which could correspond to the two invasive species, *Wisteria sinensis* and *W. floribunda*, found in the area (Figure 5).

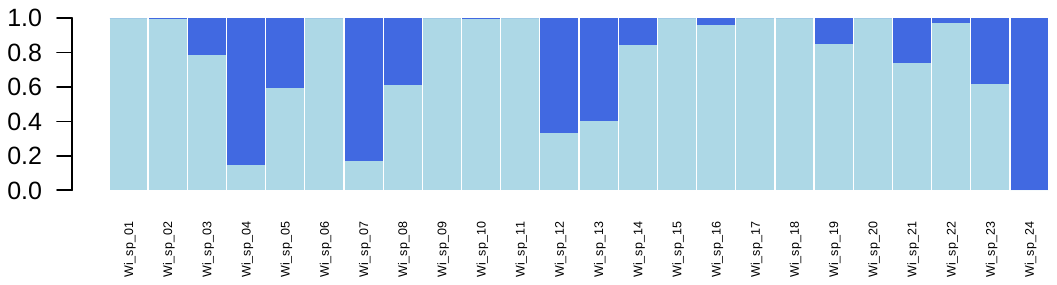

**Figure 5. Structure plot for *Wisteria* with K = 2**

Estimated from an assembly of 5,820 SNPs from 1,669 3RAD loci. Individual sample names are shown below bars.

***Project:* Rhodnius pallescens**

We included 16 samples from five populations of the kissing bug (*Rhodnius* pallescens)—a vector of the Chagas parasite—collected in Panama. We prepared libraries following the methods described above, individually amplifying each sample with 16 PCR cycles and unique iTru primers. We pooled libraries in equal amounts and size-selected for 550 bp +/- 10% on a Pippin Prep. We cleaned this product and conducteds a final 12-cycle PCR with P5/P7 primers to increase library concentration. We cleaned this product using SpeedBeads and pooled it libraries for sequencing.

We estimated F_ST_ values between 0.081 and 0.165, and our Structure analysis supported a value of *K* = 1 (Table 2; Fig. 6). These results are not unexpected, as we collected samples from a relatively small geographic area. However, localities AB130 and AB131 are located on the opposite side of the Panama Canal from the others; therefore, we expected higher levels of population differentiation between these localities and the others. We do not have specific locality information for AB234, so it is difficult to explain its relatively high F_ST_ values. However, we know that this locality is from contiguous forest, while all other samples were from forest patches, peridomestic, or pasture habitats. More data are needed to investigate the population structure of this species in Panama.

**Table 2. Pairwise F_ST_ values for *Rhodnius pallescens* from five localities**

Estimated from an assembly of 12,099 SNPs from 7,779 3RAD loci.

|  | **AB131** | **AB162** | **AB234** | **AB65** |
| --- | --- | --- | --- | --- |
| **AB130** | 0.081 | 0.094 | 0.151 | 0.093 |
| **AB131** |  | 0.095 | 0.152 | 0.096 |
| **AB162** |  |  | 0.165 | 0.093 |
| **AB234** |  |  |  | 0.147 |

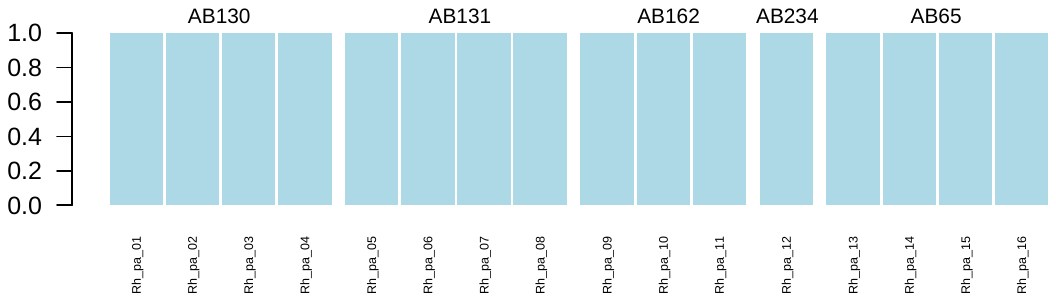

**Figure 6. Structure plot for *Rhodnius pallescens* with K = 1**

Individual sample names are shown below bars.

***Project:* Gambusia affinis**

We included 24 individuals from 11 localities in five countries (Japan, China, Taiwan, Philippines, and the native range in the United States). We prepared libraries following the methods described above, individually amplifying each sample with 12 PCR cycles and unique iTru primers. We pooled libraries in equal amounts and size-selected for 550bp +/- 10% on a Pippin Prep. We cleaned this product and conducted a final 18-cycle PCR with P5/P7 primers to increase library concentration. We quantified and pooled cleaned product with other libraries for sequencing. We used two methods to assembly data in Stacks v1.40: 1) a *de novo* assembly; and 2) a *ref_map* assembly using a reference genome. In the first, we used the *de novo* pipeline in Stacks, with the parameters as described above. In the second, we used the Burrows-Wheeler aligner (bwa) 0.7.10 to index our *Gambusia affinis* reference genome (NCBI NHOQ01000000) and align independent paired-end reads from each sample against it (bwa -mem; Li, 2013). We merged the resultant paired-reads files for each sample (using ‘cat’ Unix command). Then, we used these files as input in the Stacks v1.44 *ref_map* pipeline with the minimum number of identical reads to create a stack (-m) = 3, similar to our *de novo* approach.

We estimated F_ST_ values between 0.102 and 0.198 using the *de novo* assembly and between 0.092 and 0.183 in the *ref_map* assembly (Table 3). The largest values come from pairwise comparisons with the sample from the Philippines, which is most similar to the native population and putative source of invasive *G. affinis* throughout Asia, and less similar to other populations in Taiwan, China, and Japan than those populations are to each other. The Structure analysis supported *K* = 2 for both assemblies (Figs. 7-8), but inferred mixed ancestry varied some among assemblies.

**Table 3. Pairwise F_ST_ values for *Gambusia affinis* from five localities**

Estimated from two assemblies of 3RAD data. Values above the diagonal are calculated from the 2,140 loci recovered from *de novo* assembly and values below the diagonal are calculated from the 5,015 loci recovered from the *ref_map* assembly.

|  | **Native** | **Taiwan** | **China** | **Japan** | **Philippines** |
| --- | --- | --- | --- | --- | --- |
| **Native** |  | 0.128 | 0.126 | 0.135 | 0.102 |
| **Taiwan** | 0.118 |  | 0.124 | 0.129 | 0.170 |
| **China** | 0.115 | 0.115 |  | 0.130 | 0.198 |
| **Japan** | 0.124 | 0.117 | 0.121 |  | 0.163 |
| **Philippines** | 0.092 | 0.153 | 0.183 | 0.149 |  |

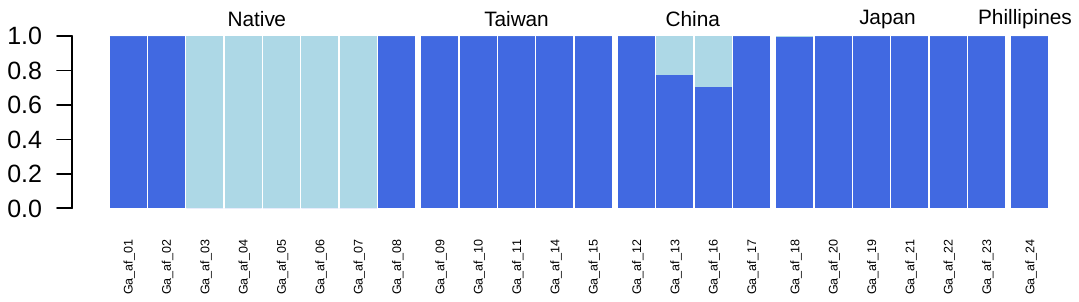

**Figure 7. Structure plot for *Gambusia affinis* with K = 2, from the *de novo* assembly**

Predefined populations are separated by large spaces and labeled above bars; individual sample names are shown below bars.

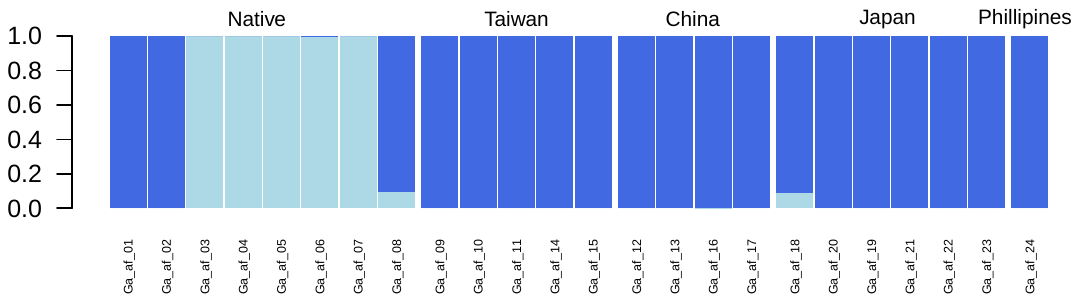

**Figure 8. Structure plot for *Gambusia affinis* with K = 2, from the *ref_map* assembly**

Predefined populations are separated by large spaces and labeled above bars; individual sample names are shown below bars.

***Projects:* Sphyrna tiburo & Sphyrna lewini**

We included 24 samples of the bonnet-head shark (*Sphyrna tiburo*) from six localities in the North Atlantic and Gulf of Mexico and 15 samples from the scalloped hammerhead shark (*S. lewini*) from seven localities in the Mexican Pacific. We assembled these data separately using the Stacks pipeline as described above.

We estimated F_ST_ values between 0.046 and 0.364 for *S. tiburo* and between 0.097 and 0.381 for *S. lewini*, and Structure results supported *K* = 1 for both species (Tables 4–5; Figs. 9–10). In *S. tiburo*, F_ST_ values were highest in pairwise comparisons between localities: 1) on the Atlantic coast of Florida and the Gulf of Mexico; and 2) offshore in Florida and the Gulf of Mexico near Mexico. Some of these results are similar to those described by previous publications (Escatel-Luna et al., 2015; Portnoy et al., 2015). In contrast, previous studies of *S. lewini* have suggested significant population structure across oceans but not along continental margins (Duncan et al. 2006). Our results likewise support the relative lack of structure in the Eastern Pacific. Results from both species encourage further research using great sample sizes to study these ecologically and commercially important sharks.

**Table 4. Pairwise F_ST_ values for *Sphyrna tiburo* from six localities**

Estimated from an assembly of 17,555 SNPs from 7,183 3RAD loci. TB: Tampa Bay, Florida; CK: Cedar Key, Florida; NAFL: North Atlantic, Florida; CAFL: Central Atlantic, Florida; TAB: Tabasco, Mexico; and Cam: Campeche, Mexico.

|  | **TB** | **CK** | **NAFL** | **CAFL** | **TAB** |
| --- | --- | --- | --- | --- | --- |
| **CAM** | 0.082 | 0.089 | 0.235 | 0.234 | 0.071 |
| **TB** |  | 0.046 | 0.073 | 0.073 | 0.051 |
| **CK** |  |  | 0.083 | 0.083 | 0.051 |
| **NAFL** |  |  |  | 0.364 | 0.079 |
| **CAFL** |  |  |  |  | 0.080 |

**Table 5. Pairwise F_ST_ values for *Sphyrna lewini* from seven localities**

Estimated from an assembly of 12,272 SNPs from 5,263 3RAD loci. BoC: Boca Camichín, Nayarit; C: Chametla, Sinaloa; LC: Las Cabras, Sinaloa; PM: Puerto Madero, Chiapas; SC: Salina Cruz, Oaxaca; and VC: Verde Camacho, Sinaloa.

|  | **C** | **LC** | **PM** | **SC** | **T** | **VC** |
| --- | --- | --- | --- | --- | --- | --- |
| **BoC** | 0.155 | 0.229 | 0.218 | 0.122 | 0.119 | 0.125 |
| **C** |  | 0.222 | 0.212 | 0.120 | 0.118 | 0.123 |
| **LC** |  |  | 0.381 | 0.155 | 0.161 | 0.167 |
| **PM** |  |  |  | 0.155 | 0.157 | 0.165 |
| **SC** |  |  |  |  | 0.097 | 0.101 |
| **T** |  |  |  |  |  | 0.097 |

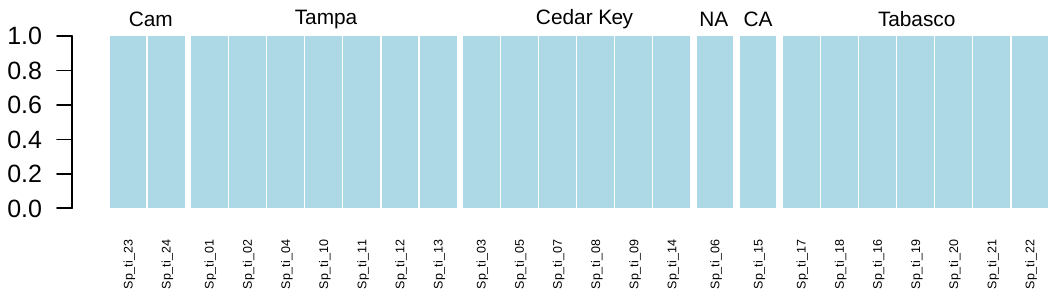

**Figure 9. Structure plot for *Sphyrna tiburo* with K = 1**

Estimated from an assembly of 17,555 SNPs from 7,183 3RAD loci. Predefined populations are separated by large spaces and labeled above bars; individual sample names are shown below bars. Cam: Campeche, Mexico; NA: North Atlantic, Florida; CA: Central Atlantic, Florida.

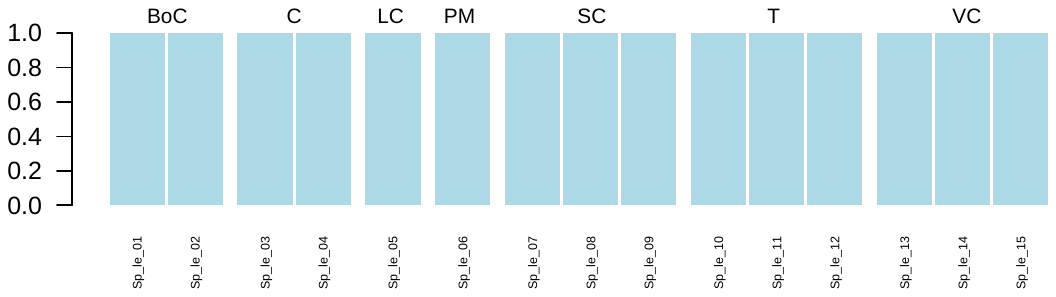

**Figure 10. Structure plot for *Sphyrna lewini* with K = 1**

Estimated from an assembly of 12,272 SNPs from 5,263 3RAD loci. Predefined populations are separated by large spaces and labeled above bars; individual sample names are shown below bars. BoC: Boca Camichín, Nayarit; C: Chametla, Sinaloa; LC: Las Cabras, Sinaloa; PM: Puerto Madero, Chiapas; SC: Salina Cruz, Oaxaca; and VC: Verde Camacho, Sinaloa

***Summary***

We demonstrate that 3RAD can be used to generate data for many variable loci across a variety of organisms. Additionally, we show that these data are useful for answering a variety of evolutionary questions. For example, when assembled differently, we recovered many homologous loci both across Kinosternidae and within the single species *S. depressus* from a single iteration of library preparations and sequencing. These data are useful for population genetic (Table 1; Fig. 2) and phylogenetic analyses (Fig. 1) across relatively deep time (ca. 55 my). We note that when species groups across deeper evolutionary time are sampled (e.g., *Eurycea* and Kinosternidae), we recover more loci per sample, but likely at the cost of reduced homology among samples.
