## Supplementary material for "Adapterama III: Quadruple-indexed, double/triple-enzyme RADseq libraries (2RAD/3RAD)": File S7

**3RAD Libraries with Molecular ID Tags Protocol**

1. Normalize all DNA samples to 20 ng/µL (quantified on Qubit), placing ≥20 µL in plates or strip tubes. Protocol will work with much less DNA, but ≥5 ng/µL is best.

2. Double Enzyme Digest Recipe (per DNA sample & unique adapter combo):

1.5 μL NEB 10x CutSmart Buffer

5.0 μL dH_2_O

0.5 µL XbaI [20.000 U/mL] (or another Read1 enzyme)

0.5 µL EcoRI-HF [20.000 U/mL] (or another Read 2 enzyme)

0.5 μL NheI [20.000 U/mL] (or another Read 1 adapter dimer cutting enzyme)

1.0 µL ds iTru NheI # adapter (5 µM) (# = A, B, … H; see Table 2)

1.0 µL ds iTru EcoRI # adapter (5 µM) (# = 1, 2, … 12; see Table 2)

5.0 µL genomic DNA (20 ng/µL; if dilute, use more volume & take out water)

3. Incubate samples at 37°C for 1 hr. in a thermal cycler, then immediately…

4. Add Ligation mix (recipe per sample):

2.0 μL dH_2_O
1.5 µL ATP (10 mM); note: rATP, NOT dATP
0.5 µL 10x Ligase Buffer (ensure components are in solution [warm it up!])
1.0 µL DNA T4 ligase [100 units/µL; dilute from 400 units/µL stock using	Diluent Buffer A]

5.0 μL Total [makes a grand total of 20 µL in each tube]

5. Incubate at 22°C for 20 min., 37°C for 10 min., 22°C for 20 min., 37°C for 10 min., 80°C for 20 min., 10°C forever.

6. Pooling step 1: Using the multi-channel pipette and changing tips each time, skloosh, then take 10 µL from each ligation and add it to a strip tube – thus from a 96-well plate, each tube in the strip will have 120 µL. Seal the plate with foil, label it well, and freeze it for potential use later.

7. Pooling step 2: Use and label two new 1.5mL tubes. Into each one, pool 60 µL from each tube of the strip from step 6. This should yield 480 µL of ligation product into each 1.5 mL tube. Label one of the 1.5mL tube well and freeze it for potential use later.

8. Split the pooled ligation products into two new 1.5 mL tubes with 240 µL of pool each (measured as 2 x 120 µL), then add 300 µL of SpeedBeads (i.e. 1.25x SpeedBeads; measured as 2 x 150 µL). Purify as normal & resuspend each in 30 µL of dH_2_O, then combine into one tube (total of 60 µL).

9. PCR recipe to add complete Illumina adapters & indexes (6 per ligation plate):

10.0 μL 5x Kapa HiFi Buffer (Kapa Biosystems, Wilmington, MA)

1.5 μL dNTP’s (10 mM stock from Kapa kit)

22.5 μL dH_2_O (to make final total volume 50 µL)

1.0 μL Kapa HiFi DNA Polymerase (1 unit/μL from Kapa kit)

5.0 µL iTru5 8N Primer (5 µM -> 0.5 µM final; note zero iTru7 primer)

10.0 μL Linker ligated DNA fragments from step 8 (placed on magnet).

1 cycle of: 98°C for 60 sec., 60°C for 30 sec., 72°C for 6 min. Hold at 15°C.

It may be easier to program this: 98°C for 40 sec.; then, 1 cycle of: 98°C for 20 sec., 60°C for 30 sec., 72°C for 60 sec.; followed by 72°C for 5 min. Hold at 15°C.

10. Pool all six reactions from above & purify with SpeedBeads (1:1.5): Add 450 µL of SpeedBeads, then adding the 6x50 µL PCR amplicons & sklooshing; purifying as normal, and resuspending in 33 µL dH_2_O.

11. PCR recipe to add complete Illumina adapters & indexes (three per ligation plate):

10.0 μL 5x Kapa HiFi Buffer

1.5 μL dNTP’s (10 mM stock from Kapa kit)

17.5 μL dH_2_O (to make final total volume 50 µL)

1.0 μL Kapa HiFi DNA Polymerase (1 unit/μL from Kapa kit)

5.0 µL P5 Primer (5µM -> 0.5 µM final)

5.0 µL iTru7 Primer(s) (5 µM -> 0.5 µM final; Note *details at end of protocol)

10.0 μL PCR product from step 10 (placed on magnet).

Cycle: 98°C for 2 minutes.; then, 6 cycles of: 98°C for 20 sec., 60°C for 15 sec., 72°C for 30 sec.; followed by 72°C for 5 min. Hold at 15°C. [Note: Adjust the number of cycles based on the amount of input DNA, more cycles if less input DNA was used.]

12. Pool all three reactions from above & purify with SpeedBeads (Thermo-Scientific, Waltham, MA, USA) to a 1:1.5 ratio, DNA: SpeedBeads. Add 225 µL of SpeedBeads, then add the 3x50 µL PCR amplicons & skloosh; purify as normal, and resuspend in 60 µL dH_2_O, then place on magnet and pull off all liquid (~55 µL), transferring the clean solution to a new labeled tube and leaving the beads behind.

13. Run 5µL on agarose gel to ensure each sample worked.

14. Quantify with Qubit, normalize, pool, SpeedBeads (1:1.25, DNA: SpeedBeads), and size select on Pippin Prep (525 bp +/- 10%). [change size as necessary to avoid bright bands; we do *not* know that choosing a narrow size-range is best]

15. Quantify with Qubit (and qPCR), mix in appropriate proportions with other libraries, and send off to HiSeq or NextSeq: Use TruSeq sequencing primers & do dual (8nt) index reads!

Special notes & considerations:

*How many iTru7 primers to use (step 11) will vary depending on how many plates will be pooled together for sequencing. If you are running:

1) ≥12 plates in (or no single plate you produce will take up ≥10% of) a HiSeq lane or NextSeq/MiSeq/MiniSeq/NovaSeq run, then use a single iTru7 primer for all 3 PCR reactions. This will make your life easiest.
2) Only a few plates in a HiSeq lane or NextSeq/MiSeq/MiniSeq/NovaSeq run, then you will want to use multiple iTru7s. In general, you need base diversity of the indexes to be complex for them to sequence correctly. Read the note for a single plate of samples and use the index diversity calculator for your scenario.
3) A single plate of samples in a HiSeq lane or NextSeq/MiSeq/MiniSeq/NovaSeq run, then either use 3 different iTru7’s that you verify can sequence together correctly (i.e., by using the index diversity calculator), or by using pools of iTru7 primers to yield a full set of 12 (we do not know if it is best to use 3 different groups of 4 or the same group of 12 in all 3 PCRs). It may seem stupid to do this because you will demultiplex then repool (concatenate) the reads back together, but this is necessary to generate usable iTru7 index data, which then behave appropriately in *stacks* and *ipyrad*.
